# Novel inhibitors against COVID-19 main protease suppressed viral infection

**DOI:** 10.1101/2022.11.05.515305

**Authors:** Vijayan Ramachandran, Yanyun Liu, Qianying He, Andrew Tang, Patrick Ronaldson, Dominik Schenten, Rui Chang

## Abstract

Severe acute respiratory syndrome coronavirus 2 (SARS-CoV-2), the etiologic agent of COVID-19, can cause severe disease with high mortality rates, especially among older and vulnerable populations. Despite the recent success of vaccines and approval of first-generation anti-viral inhibitor against SARS-CoV-2, an expanded arsenal of anti-viral compounds that limit viral replication and ameliorate disease severity is still urgently needed in light of the continued emergence of viral variants of concern (VOC). The main protease (Mpro) of SARS-CoV-2 is the major non-structural protein required for the processing of viral polypeptides encoded by the open reading frame 1 (ORF1) and ultimately replication. Structural conservation of Mpro among SARS-CoV-2 variants make this protein an attractive target for the anti-viral inhibition by small molecules. Here, we developed a structure-based *in*-*silico* screening of approximately 11 million compounds in ZINC15 database inhibiting Mpro, which prioritized 9 lead compounds for the subsequent *in vitro* validation in SARS-CoV-2 replication assays using both Vero and Calu-3 cells. We validated three of these compounds significantly inhibited SARS-CoV-2 replication in the micromolar range. In summary, our study identified novel small-molecules significantly suppressed infection and replication of SARS-CoV-2 in human cells.

## Introduction

Coronaviruses comprise a large family of positive single stranded RNA viruses that cause respiratory, gastrointestinal, and neurological diseases in humans and other animals ^1,2^. Severe acute respiratory syndrome coronavirus 2 (SARS-CoV-2) ^3–6^, the etiological agent of COVID-19 and its ever-increasing evolutionary variants, has become a global health emergency with an urgent need for novel therapeutic strategies to combat the disease. Despite the remarkable and rapid success of vaccines against SARS-CoV-2^7^ in the U.S. and other developed countries, significant infection risk remains among unvaccinated people, immunocompromised or otherwise vulnerable individuals forming a substantial reservoir to support viral spread, which make small-molecule inhibitors of SARS-CoV-2 replication urgently needed. In addition, vaccines mitigate but do not eliminate the likelihood of severe disease. Finally, the emergence of SARS-CoV-2 variants of concern (VOCs), such as the latest strain Omicron, which is highly contagious with increased likelihood to escape vaccine-derived immune surveillance, have raised concerns about the efficacy of the current vaccines, thus illustrating the importance of a wide arsenal of tools to combat the evolving current SARS-CoV-2 strains or novel future coronaviruses altogether.

The SARS-CoV-2 genome encodes several structural proteins including the membrane (M), envelope (E), and spike (S) proteins as well as multiple non-structural proteins that are necessary for viral replication or the manipulation of the host immune response ^8–10^. The main protease (Mpro, also known as 3CLPro) is a cysteine protease that is critical for the cleavage of two polypeptide chains encoded by the overlapping open reading frames ORF1a and ORF1b into functional proteins ^11,12^. Among these proteins is the essential RNA polymerase RdRp that is responsible for the replication of viral genome and whose activity is severely compromised without prior proteolytic cleavage by Mpro. In addition to the processing of viral proteins necessary for the viral replication machinery, Mpro has also been suggested to interfere with the induction of cellular type I and type III interferon (IFN) and proinflammatory cytokine responses, either directly through the proteolytic cleavage of members of the IFN signaling cascade or indirectly by promoting the processing of other viral proteins that themselves interfere with IFN signaling^13–15^. The pharmacological inhibition of Mpro may therefore also limit viral replication by inducing a type I and type III IFN-dependent anti-viral state of the host cells.

To identify novel inhibitors of Mpro, we developed an *in-silico* pipeline to screen compounds in the ZINC15 database against Mpro and prioritized 9 lead compounds. We then validated the function of these lead compounds by using replication assays with SARS-CoV-2 in both rhesus monkey kidney-derived Vero cells and human lung-derived Calu-3 cells and identified three novel compounds that can significantly suppress the replication of SARS-CoV-2 by interfering with Mpro, therefore these compounds serve as starting point for further drug development.

## Results

### Mpro is conserved among SARS-CoV-2 variants of concern

The frequent occurrence of mutations in the viral spike (S) protein among SARS-CoV-2 VOCs suggest that the S protein of SARS-CoV-2 remains under evolutionary pressure to adapt to the human ACE2 receptor. Mpro may be less sensitive to such selective pressure as it has a substrate specificity for viral proteins with unique glutamate-containing cleavage sites that are distinct from the sites used by known human proteases ^16^. To evaluate whether Mpro is indeed structurally conserved among known SARS-CoV-2 strains, we compared the consensus sequence of original North-American WA1 strain to a range of SARS-CoV-2 variants of concern, including B.1.1.7(Alpha), B.1.351(Beta), B.1.427 & B.1.429 (Epsilon), B.1.525 (Eta), B.1526 (Iota), B.1.526.1, B.1.617.1 (Kappa), B.1.617.2 (Delta), P.1 (Gamma), P.2 (Zeta) and B.1.1.529 (Omicron), as well as the more distantly related human coronaviruses SARS-CoV and MERS. We found that there are only three substitutions (K90R, L205V, P132H) in Mpro across all VOCs of SARS-CoV-2 (Online Method, Figure 1A, Supplementary Figure 1, Supplementary Table 1). The recently determined X-ray crystal structure of Mpro (PDB ID: 6W63) reveals four distinct regions of the main protease protomer, namely Domain I (residue 10-99) and Domain II (residue 100-184) that form antiparallel β-barrels, followed by a long connecting region (residues 185-200), and finally Domain III (residues 201-303) that forms a cluster of helices ^17^. The substrate binding site is located in the cleft between domain I and domain II ^17^. Compared to WA1, we found only three mutations in Mpro, namely a K90R mutation in the B1.351 (Beta) strain and a L205V mutation in the P2 (Zeta) strain, and P132H mutation in B.1.1.529 (Omicron), none of the mutated residues were part of the substrate binding site (Figure 1A). In contrast, the mutation rate was significantly higher in S protein with 78 mutations (substitution, deletion, and insertion) (Online Method, Figure 1B, Supplementary Figure 2, Supplementary Table 1) ^18^. Together, this comparison suggests that the substrate binding site of Mpro is more conserved among all known human SARS coronaviruses, thus making this an ideal anti-viral drug target.

**Figure 1.**
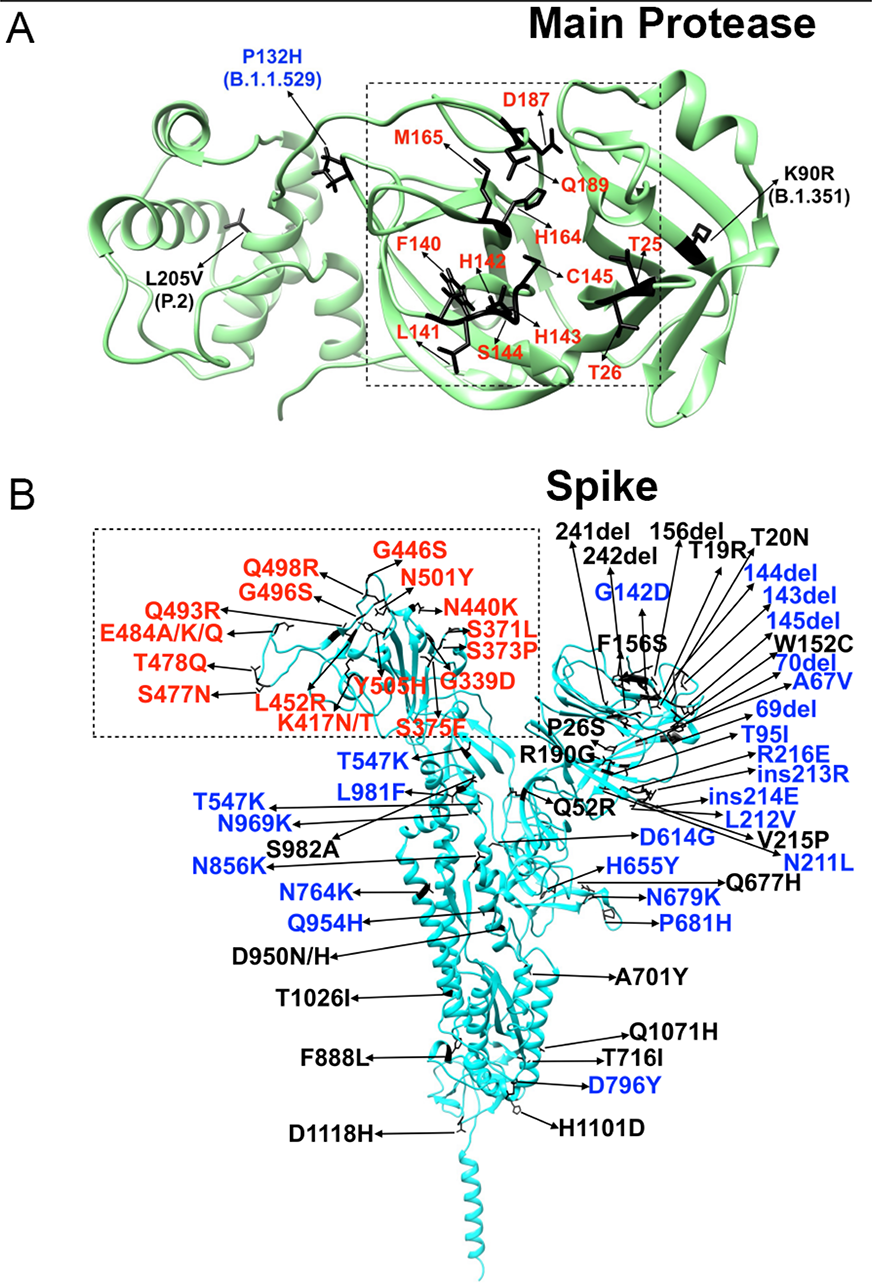
The structure of Mpro and Spike protein across SARS-CoV-2 variants. (A) The active site residues of Mpro are highlighted in red color. Omicron-specific mutation, P132H (B. 1.1529) of Mpro protein are highlighted in blue. Mutations of Mpro protein (K90R, L205V) from the B.1.351 and P.2 variants are highlighted in black. (B) The active site residues of Spike protein are highlighted in red. Omicron-specific mutations of Spike protein are highlighted in blue. Mutations of Spike protein from all the other VOCs are highlighted in black.

### Integrative *in-silico* screening identified novel candidate inhibitors of Mpro

Structure-based virtual screening is a fast and powerful method for lead compound discovery. We developed an integrative approach for *in-silico* screening and prioritizing (Figure 2, Online Method) compounds targeting Mpro of SARS-CoV-2. In step 1, we downloaded 11 million Drug-Like In-Stock 3D small molecules from ZINC15 database^19^ and the X-ray crystal structure of Mpro from RCSB Protein Data Bank. In step 2, we prepared the X-ray crystal structure of Mpro with the Protein Preparation Wizard in Maestro from Schrödinger and prepared ligand by removing compounds with reactive functional groups to obtain 10.4 M compounds. In step 3, we defined the binding pockets of Mpro based on the reported inhibitor X77 (https://www.rcsb.org/structure/6W63). In step 4, we performed virtual screening with Schrödinger ^20^ to select top-500 lead compounds(Online Method). In step 5, we prioritize the top-ranked 500 lead compounds by integrating two complementary strategies (Online Method), which lead to the selection of total 9 lead compounds (Supplementary Table 2) for *in-vitro* testing.

**Figure 2.**
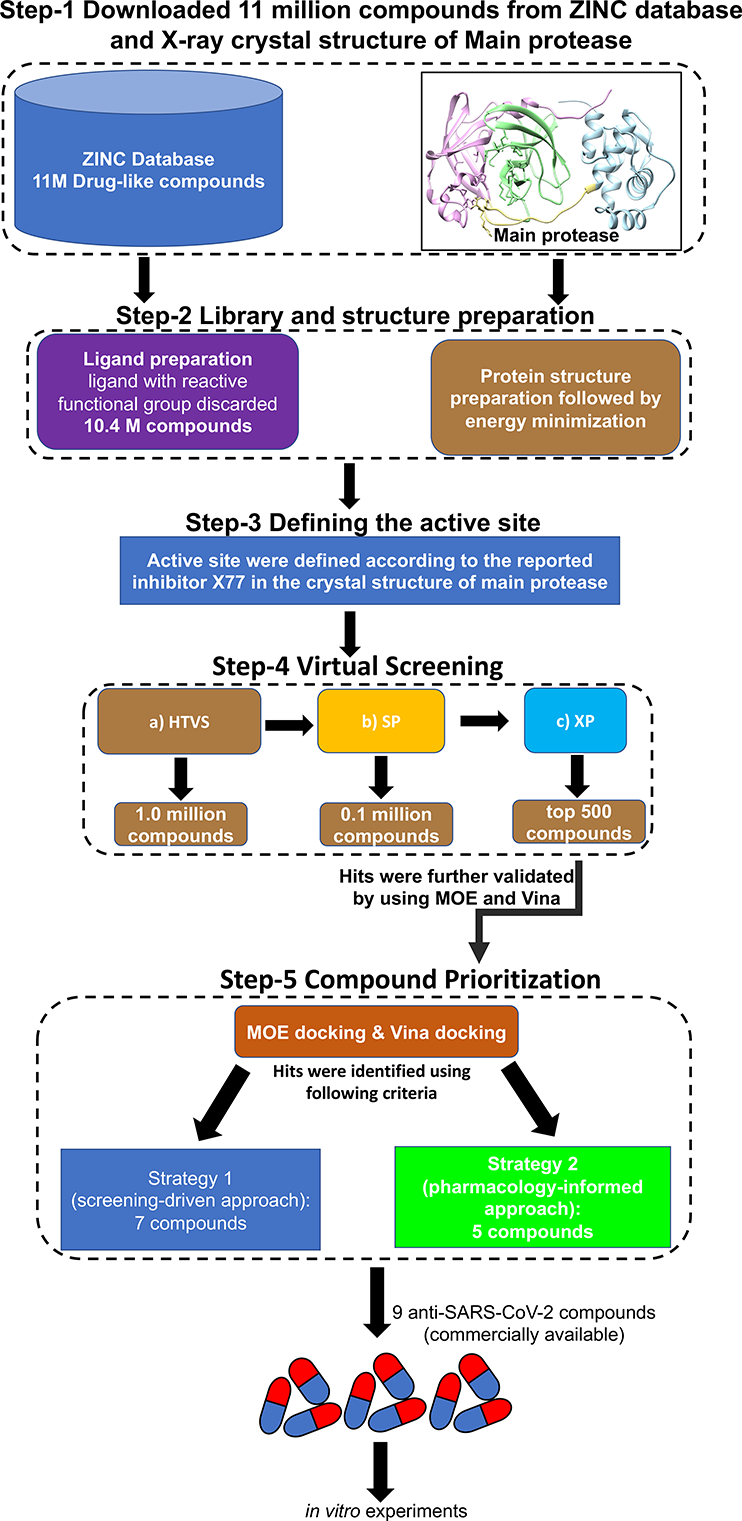
*In-silico* screening workflow of SARS-CoV-2 Mpro inhibitor. Workflow for the identification of potential small molecular inhibitors for SARS-CoV-2 Mpro. The structure-based virtual screening was carried out using Schrödinger, MOE, Vina. After virtual screening, the top-ranked 500 lead compounds were prioritized based on the Schrödinger Glide score and further independently prioritized using two strategies that ultimately led to total of 9 lead molecules for *in-vitro* testing.

The identified 9 lead compounds showed higher binding affinity with Mpro (Supplementary Table 2). From the molecular interaction analysis of docked complexes, we observed that all the 9 lead compounds show hydrogen bonding interactions and other potential hydrophobic or hydrophilic interactions with Mpro. All the 9 lead compounds were bound to the same binding site. Residue Leu141, Asn142, Gly143, Glu166, and Gln189 were the common interacting residues between Mpro and 9 lead compounds, which suggested the crucial role of these residues in stabilizing the Mpro-ligand complex. Importantly, to demonstrate our 9 lead molecules are robust against VOCs of SARS-CoV-2, we built Mpro structures with K90R mutation in the B1.351 (Beta), L205V mutation in the P2 (Zeta), and P132H mutation in B.1.1.529 (Omicron), by using UCSF Chimera^21^ followed by the Protein Preparation Wizard in the Schrödinger package and validated all compounds by docking them to these three mutant structures of Mpro. The docking results showed very similar binding affinities of these lead compounds to the mutant structures compared to wild-type Mpro suggesting that these lead compounds have the potential for the robust inhibition of the currently known VOCs of SARS-CoV-2.

### *In vitro* validation of inhibitors of SARS-CoV-2 infection

We tested the ability of the selected 9 lead compounds to suppress SARS-CoV-2 infection in two experimental settings. Initially, we infected Vero cells with SARS-CoV-2 immediately after the addition of increasing concentrations of the lead compounds and measured the number of infected cells by viral reduction plaque assay 4 days later. Cells infected in the absence of the compounds served as positive control, whereas uninfected cells cultured in the presence of increasing concentrations of the compounds served as control for cytotoxicity. We observed a significant decrease in the number of infected cells compared to positive control for three of the 9 lead compounds, i.e. PATH-6, PATH-7, and PATH-8. We measured the cytopathic effect (CPE) on Vero cells 4 days after infection with SARS-CoV-2 in the presence of increasing concentrations 0.001 to 100 μM of the three compounds, starting at the time of infection (Figure 3). In parallel, we also determined the cytotoxicity of these compounds over the same dose range. These experiments revealed that PATH-6, PATH-7, and PATH-8 inhibited virus-induced CPE with an EC50 between 1.57 – 9.3 μM (Figure 3A-C). Importantly, PATH-6 showed no sign of cytotoxicity (Figure 3A) and PATH-7 showed no sign of cytotoxicity within 10 μM (Figure 3B). Overall, these experiments confirmed 3 out of the total 9 lead compounds significantly suppressed SARS-CoV-2 infection *in vitro* and especially PATH-6 is a promising compound for further investigation.

**Figure 3.**
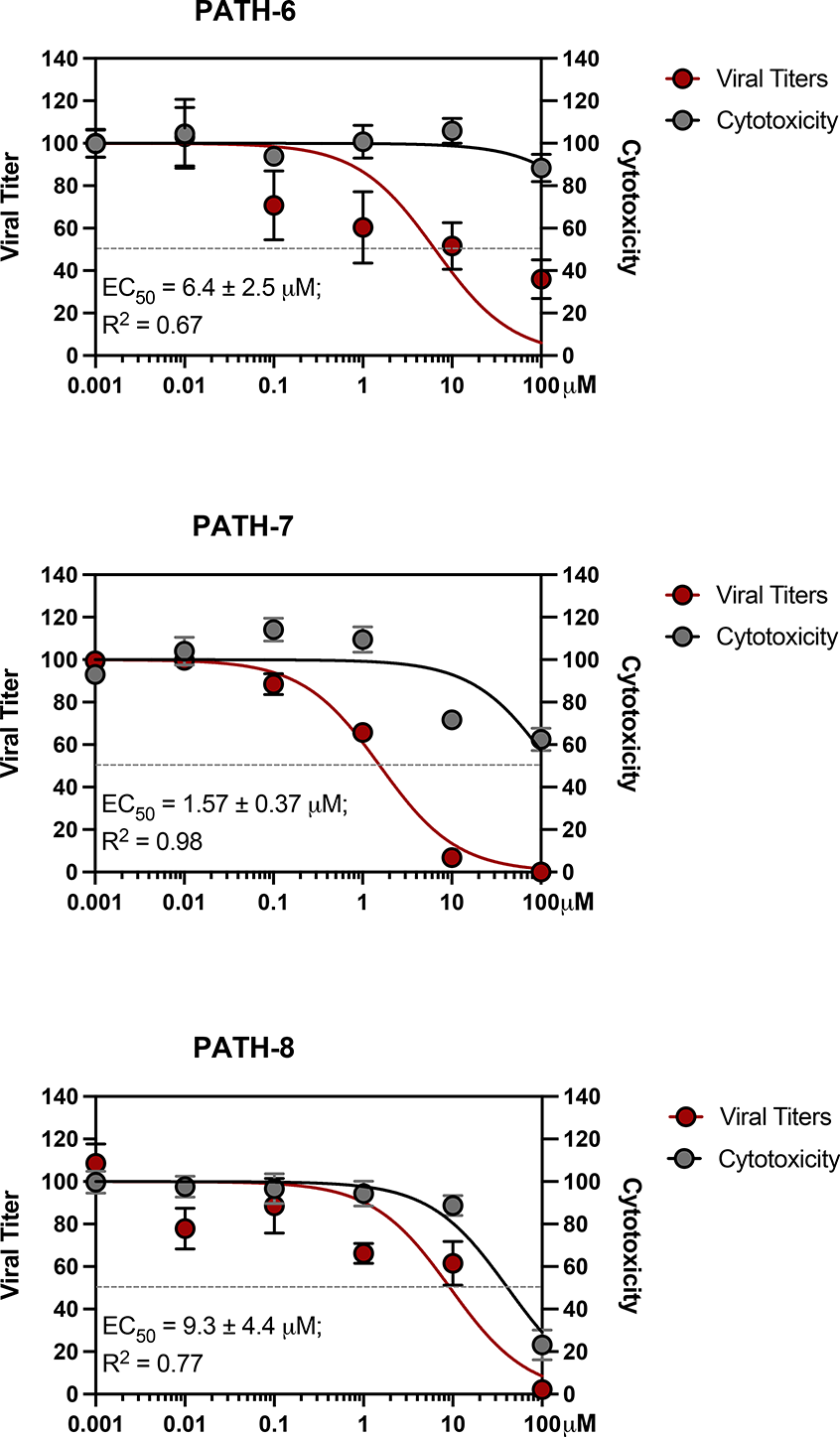
CPE and cytotoxicity in Vero cells treated with putative SARS-CoV-2 Mpro inhibitors. **(A-C)** Vero cells were incubated with the P-gp inhibitor CP-100356 and infected with SARS-CoV-2 in the presence of increasing concentrations of putative Mpro inhibitors. CPE (red line) and cytotoxicity (black line) of the inhibitors was determined 3 days later and expressed as fraction of adherent cells in infected samples relative to uninfected controls (CPE) or untreated cells (cytotoxicity). The amount of adherent cells was quantified by the spectrophotometrical measurement of crystal violet staining. Shown are the results of (A) PATH-6, (B) PATH-7, (C) PATH-8. Each experiment was set-up in triplicates and independently repeated three times. The EC50 of viral inhibition and the 95% confidence interval is indicated in each graph.

## Discussion

Mpro, the main protease of SARS-CoV-2, plays a central role in the cleavage of the ORF1-encoded pp1a and pp1ab polypeptides to produce active viral proteins, including the RNA polymerase RdRP. Pharmacological inhibition of Mpro therefore likely inhibits SARS-CoV-2 infection directly by preventing the replication of viral RNA genomes. Across currently 12 VOCs of SARS-CoV-2, the sequence of Mpro is significantly more conserved than the sequence of Spike protein, with only 3 mutations outside its known binding pocket. Therefore, Mpro represent an attractive drug target to interfere with viral replication. Recent studies have reported the possible inhibitors against Mpro of SARS-CoV-2^22–26^. Indeed, the recent approval of Paxlovid (PF-07321332), a derivative of the Mpro inhibitor GC376, demonstrated this target^7^. While the emergence of the first clinical SARS-CoV-2 Mpro inhibitor is encouraging, it is also clear that the continued expansion of the arsenal of anti-viral drugs is highly desirable.

In the present study, we developed an *in-silico* screening pipeline which integrated the structure-based docking with pharmacokinetics prediction to prioritize approximately 11 million compounds for their ability to bind to Mpro and to block its enzymatic activity in the process of viral replication. To cross-check the binding affinities of the experimentally validated compounds predicted by our *in-silico* screening pipeline, we examined the binding mode of these compounds; PATH-6 occupied the binding pocket with a Glide docking score of −11.79 kcal/mol, Vina score of −8.73 kcal/mol, and MOE score of −14.81 kcal/mol. It forms seven hydrogen bonds with residues of Gly143, His164, Glu166, Thr190 and Gln192. PATH-7 fitted in the binding pocket with a Glide docking score of −8.99kcal/mol, Vina score of −7.77 kcal/mol, and MOE score of −6.86 kcal/mol. There are three hydrogen bonds formed between this compound and residue Ser144, Cys145, and Gln189. PATH-8 located in the binding site with a Glide docking score of −9.00 kcal/mol, Vina score of −9.97 kcal/mol, and MOE score of −7.6 kcal/mol. Four hydrogen bonds were formed between the compound and residues Cys44, Thr190, and Gln192. The hydrogen bond, van der Waals, hydrophobic and Pi-Pi interactions between the validated three compounds and SARS-CoV-2 Mpro mainly occurred at the catalytic pocket with strong bonding to His41 and Cys145, indicating these residues may be particularly responsible for inhibiting SARS-CoV-2 Mpro.

To test their inhibitory activity of SARS-CoV-2, we performed two rounds of *in vitro* experiments. The initial *in vitro* infection of cells with SARS-CoV-2 identified three out of the 9 lead compounds (PATH-6, PATH-7 and PATH-8) as potentially inhibitors of SARS-CoV-2 infection. We further fully confirmed the inhibition of SARS-CoV-2 infection for these compounds using a diverse set of assays in two distinct cell lines, namely African green monkey Vero cells and human lung Calu-3 cells.

In summary, our study identified 3 novel compounds, i.e. PATH-6, PATH-7, and PATH-8, that not only directly inhibit viral replication by binding to Mpro, but also released the suppression of cellular anti-viral immune response by the virus. Hence, our antiviral compounds can be further optimized and added to the existing reservoir of the COVID-19 Mpro inhibitors for future drug development against SARS-CoV-2 and related coronaviruses.

## Materials and methods

### Sequence alignment of Spike and 3C-like proteinase and its variants

The protein sequences of Spike glycoprotein variants and 3C-like proteinase variants of SARS-CoV-2 isolates were retrieved from NCBI https://www.ncbi.nlm.nih.gov/datasets/coronavirus/proteins/). The Spike protein, 3C-like proteinase and its variants sequences were extracted using our UNIX script. To determine the level of the conservancy, multiple sequence alignment (MSA) was performed for the sequences using the BioEdit-ClustalW multiple alignment program (http://www.mbio.ncsu.edu/BioEdit/bioedit.html). After multiple alignment for all the download sequences for each variant, we created a consensus sequence. Next, we used Clustal Omega (https://www.ebi.ac.uk/Tools/msa/clustalo/) to align consensus sequence across all the VOCs of SARS-CoV-2, SARS-CoV, and MERS-CoV.

### Virtual screening

To identify the potential small molecular inhibitors for SARDS-CoV-2 Mpro, the structure-based virtual screening was carried out by Virtual Screening Workflow (VSW) from Schrödinger suites^20^. A total around 11 million Drug-Like In-Stock 3D small molecules obtained from ZINC15 database^19^ were performed through VSW. Mpro structure was prepared using Protein Preparation Wizard in Maestro from Schrödinger^29^. Hydrogens were added to the protein and bond orders were assigned. All hydrogen-bonding networks were optimized, and the ionization states were assigned at pH 7.0. OPLS3e force field was used for restrained minimization. The docking grid was generated with the Receptor Grid Generation tool from Maestro. The default van der Waals scaling factor of 1.0 and partial charge cutoff at 0.25 was applied^30^. The center and the size of the grid box were defined according to the position of the published inhibitor X77 in the crystal structure of Mpro (PDB ID: 6W63). To dock ligands with suitable pharmacological property, the small molecules were prefiltered by removing ligands with reactive functional groups. Around 10.4 million compounds were obtained. Virtual screening was carried out in three sequential steps, namely (a) Glide high throughput virtual screening (HTVS) docking; After HTVS docking, (b) Glide standard precision (SP) docking, and finally (c) Glide extra precision (XP) docking. At each step, the top 10% of the compounds were advanced to the next step. Finally, according to the Glide score and protein-ligand interactions, top 500 lead compounds were selected for further evaluations.

### Auto Dock Vina and MOE docking

Different docking methods, including Auto Dock Vina docking and MOE docking, were used to predict the binding affinity of the top 500 lead compounds which were prioritized based on previous Glide score against Mpro. Auto Dock Vina is a widely used open-source docking program^31^. The Mpro and structures of the top 500 lead compounds were prepared with Auto Dock Tools ^32^. All hydrogens were added and Gasteiger charges were assigned. The center and the size of the grid box were defined according to the position of the published inhibitor X77 (https://www.rcsb.org/structure/6W63). The level of exhaustiveness was set to 8. The Auto Dock Vina docking score was used to rank the binding affinity of different docking poses. The docking calculations were repeated 3 times with different random seeds.

Next, we employed MOE docking ^33,34^ to calculate the binding affinity of the top 500 lead compounds. The protein was kept as rigid, and a maximum of 30 conformations for each ligand was tested, using the default parameters of MOE using Triangle Matcher placement. The top ranked conformations of lead molecules were stored. On the basis of MOE scoring (London dG), binding free energy calculation in the S field was scored, as the London dG is a scoring function that estimates the free energy of binding of the ligand for a given pose. For all scoring functions, lower scores indicate more favorable poses.

### Toxicity Prediction

The *in-silico* toxicity properties were predicted by Data Warrior ^35^. Data Warrior was used to predict the molecular weight (MW), mutagenicity, tumorigenicity, and irritant properties as well as pharmacokinetic properties, Topological Molecular Surface area (TPSA), partition coefficient (log*P*) for the identified top 500 molecules. Toxicity risks were predicted from precompiled lists of fragments using an algorithm that gives rise to toxicity alerts in case they encounter the structure in evaluation ^36^.

### Top 500 lead compounds prioritization

To prioritize robust compounds for experiment validation, we developed an ensemble of two complementary strategies. In the first strategy, we employed a *screening-driven* approach to prioritize robust lead compounds. We first ranked the top 500 lead compounds based on the Glide score, then we focus on the top 50 lead compounds. Next, we rank these top 50 lead compounds according to the average (priority score) of the individual Glide, Vina and MOE docking scores rank. Then, we calculated the binding mode clustering by Schrödinger and structure similarity clustering by Data Warrior. Among the top 50 compounds, we identified 10 binding mode clusters and 23 structure similarity clusters. We removed 6 compounds based on the predicted toxic pharmacological properties by Data Warrior. For the remaining 44 compounds, we first selected the top compound with the highest priority score within each binding mode cluster to obtain total 9 compounds. Next, we examined the 9 compounds against their structure similarity and selected the top compound per structure similarity cluster with highest priority score. Note that 3 of the 9 compounds belong to the same similarity cluster, so the total number of selected compounds is 7. Finally, 4 out of the final selected 7 compounds are commercially available and are used for the subsequent *in-vitro* infection experiments.

In the second strategy, we employed a pharmacology-informed approach to prioritize the top 500 compounds by integrating the docking score (Glide score) with molecular weight (MW) mutagenicity, tumorigenicity, and irritant properties as well as pharmacokinetic properties, the total polar surface area (TPSA) ^37^ and the partition coefficient (LogP) ^38^ of these compounds. First, we removed 322 compounds with molecular weight greater than 500 Daltons resulting in 178 compounds. Second, we used the same toxicity criteria as above to remove 27 potentially toxic compounds. Third, we ranked the remaining 151 compounds according to their Glide score which ranges from −10.67 to −8.87 and we divided the scores into two bins: [−10.67, −9] and [−9, −8]. We selected the top compound from each bin. Fourth, since natural compounds are more accessible, can have anti-viral effects against SARS-CoV-2^39–44^, and may help to accelerate drug development, we focused on prioritizing natural compounds for the remaining 149 compounds, of which 16 are natural compounds according to ZINC database classification ^19^. Next, since drug-like molecules should be water-soluble to reach target tissues and enter cells through passive mechanisms such as the diffusion through cellular membranes. The ideal distribution coefficient for the tested compounds should therefore be neither too lipophilic nor too hydrophilic. Such pharmacological properties determines the good absorption and distribution *in vivo* and guide the translation of chemical inhibitors or viral replication into successful drugs for patients ^45^. To select drug-like natural compounds, we set TPSA values between 118 and 148 and LogP values between 2 and 4 following previous published practice ^37,38^, which resulted in 3 out of the 16 natural compounds. In total, we prioritized 5 compounds by pharmacology-informed approach which were all commercially available for experimental testing.

In summary, we selected 7 compounds by screening-driven approach, out of which 4 compounds are commercially available, and 5 compounds by pharmacology-informed approach, all of which are commercially available. Therefore, total 9 compounds were ordered for experimental validations.

#### Cells

Vero and Calu-3 cells were obtained from the American Type Culture Collection (ATCC) and cultured according to the recommendations provided by the ATCC. The cells were routinely monitored for the absence of mycoplasma infection.

#### Compounds

Potential inhibitors of Mpro identified in the virtual screens were obtained from MolPort and eMolecules and dissolved at 10 mM in PBS or PBS + 10% DMSO. The compounds were further diluted >100-fold in tissue culture medium containing 5% FCS to obtain working concentrations for the viral replication assays.

#### Virus production

SARS-CoV-2 strain WA1 and West Nile virus (WNV) strain NY99 were obtained from BEI Resources and propagated in Vero cells. The cells were infected with SARS-CoV-2 at an MOI of 0.005 and with WNV at an MOI of 0.01. After 48 hr (SARS-CoV-2) or 72 hr (WNV) of culture, the cells were harvested with a cell scraper and spun and together with the culture medium at 3000 rpm for 10 min. Supernatants were set aside while the resuspended cell pellets were treated with a Dounce homogenizer and subjected to two freeze-thaw cycles before combined with the original supernatants. Following an additional centrifugation step, supernatants were aliquoted, frozen, and subsequently titered in serial dilutions by viral plaque assay. All work with SARS-CoV-2 and WNV was performed under BSL3 conditions in a facility with negative pressure and PPE that included Tyvek suits and N95 masks for respiratory protection.

#### Viral plaque and foci assays

The number of infectious SARS-CoV-2 virions was quantified by viral plaque assay. To this end, Vero cells were incubated with SARS-CoV-2 for 2 hr and subsequently overlaid with 1% methylcellulose in culture medium. After 3-4 days, the cells were fixed in 10% formalin for 30 min, washed under tap water, and stained with crystal violet. The number of plaques corresponding to infections of individual cells by single virions was counted on a light table. The quantification of infectious WNV virions was performed similarly with the exception that the number of infected cell foci was determined by intracellular staining using a biotinylated anti-WNV-E antibody (clone E16), followed by an HRP-labeled anti-streptavidin antibody. HRP activity was detected with KPL Trueblue substrate (SeraCare).

#### Cytopathic effect and cytotoxicity measurements and other assays to determine drug activity

Viral replication of SARS-CoV-2 in the presence of Mpro inhibitors was measured in Vero or Calu-3 cells in three ways. In the first approach, Vero or Calu-3 cells were infected with SARS-CoV-2 at an MOI of 0.01in the presence of indicated concentrations of Mpro inhibitors and, if applicable, with 1.5 μM of the P-gp inhibitor CP-100356. The virus-induced cytopathic effect was measured by determining the fraction of formalin-fixed adherent cells that remained after 3 days (Vero cells) or 4 days (Calu-3 cells). To this end, the cells were stained with crystal violet, PBS-washed, and air-dried. Following resuspension in methanol, crystal violet staining was measured in the Spectrophometer at OD594. Cytotoxicity of the compounds was measured in parallel by the staining of uninfected cells incubated with the Mpro. In a second approach, drug activity was determined in Vero cells directly by viral plaque assay, using between 100-500 Pfu/well of virus and indicated concentrations of Mpro inhibitors. In a third approach, Vero cells were infected with SARS-CoV-2 or WNV at an MOI of 0.01 in the presence of 100 mM of Mpro inhibitors for 2-3 days. Viral replication was measured indirectly at indicated time points by quantifying the titers of infectious virions in the supernatants with viral plaque or foci assays in the absence of the compounds. Viral replication in Calu-3 cells in the presence of indicated concentrations of compounds was determined similarly.

#### RT-qPCR

RNA was isolated from infected cells, reverse-transcribed with random hexamers, and subsequently amplified using the SYBR mix on a Quant3 Real-Time PCR cycler.

## Acknowledgements

We would like to thank Jennifer Uhrlaub and the University of Arizona Keating BIO5 BSL3 facility for expert technical advice, logistical help, and overall facility management. We also like to thank Dr. Wei Wang at the University of Arizona for scientific feedback and critical reading of the manuscript. Dr. Rui Chang is the founder of INTelico Therapeutics LLC and co-founder of PATH BIOTECH LLC. Dr. Patrick Ronaldson is the co-founder of PATH Biotech LLC. This study is not supported by any company.

---

**Supplementary Figure 1. Sequence comparison of Mpro across SARS-CoV-2 variants and other human β-coronaviruses**. The locations of non-synonymous mutations resulting in a lysine to arginine mutation in the SARS-CoV-2 B.1.351 (Beta) variant, a leucine to valine mutation in the SARS-CoV-2 P.2 (Zeta) variant, and proline to histidine mutation in the SARS-CoV-2 B.1.1.529 (Omicron) variant are shown in red box.

**Supplementary Figure 2. Sequence comparison of Spike protein across SARS-CoV-2 variants and other human β-coronaviruses**. Mutations of Spike protein across all VOCs of SARS-CoV-2 are highlighted in red box. The mutation rate was significantly higher in Spike protein with 78 mutations (substitution, deletion and insertion).

**Supplementary Table 1**: Mutations of Mpro and Spike protein across SARS-CoV-2 VOCs.

**Supplementary Table 2**: Summary of in-silico screening results for the prioritized 9 lead inhibitors of Mpro of SARS-CoV-2.

## Notes

### Competing Interest Statement

The authors have declared no competing interest.

