## Supplementary material for "Novel inhibitors against COVID-19 main protease suppressed viral infection": Figure S1

```
SARS                 SGFRKMAFPSGKVEGCMVQVTCGTTTLNGLWLDDTVYCPRHVICTAEDMLNPNYEDLLIR   60
MERS                 SGLVKMSHPSGDVEACMVQVTCGSMTLNGLWLDNTVWCPRHVMCPADQLSDPNYDALLIS   60
WA1(COVID-19)        SGFRKMAFPSGKVEGCMVQVTCGTTTLNGLWLDDVVYCPRHVICTSEDMLNPNYEDLLIR   60
B.1.1.7(Alpha)       SGFRKMAFPSGKVEGCMVQVTCGTTTLNGLWLDDVVYCPRHVICTSEDMLNPNYEDLLIR   60
B.1.351(Beta)        SGFRKMAFPSGKVEGCMVQVTCGTTTLNGLWLDDVVYCPRHVICTSEDMLNPNYEDLLIR   60
B.1.427(Epsilon)     SGFRKMAFPSGKVEGCMVQVTCGTTTLNGLWLDDVVYCPRHVICTSEDMLNPNYEDLLIR   60
B.1.429(Epsilon)     SGFRKMAFPSGKVEGCMVQVTCGTTTLNGLWLDDVVYCPRHVICTSEDMLNPNYEDLLIR   60
B.1.525(Eta)         SGFRKMAFPSGKVEGCMVQVTCGTTTLNGLWLDDVVYCPRHVICTSEDMLNPNYEDLLIR   60
B.1.526(Iota)        SGFRKMAFPSGKVEGCMVQVTCGTTTLNGLWLDDVVYCPRHVICTSEDMLNPNYEDLLIR   60
B.1.526.1            SGFRKMAFPSGKVEGCMVQVTCGTTTLNGLWLDDVVYCPRHVICTSEDMLNPNYEDLLIR   60
B.1.617.1(Kappa)     SGFRKMAFPSGKVEGCMVQVTCGTTTLNGLWLDDVVYCPRHVICTSEDMLNPNYEDLLIR   60
B.1.617.2(Delta)     SGFRKMAFPSGKVEGCMVQVTCGTTTLNGLWLDDVVYCPRHVICTSEDMLNPNYEDLLIR   60
P.1(Gamma)           SGFRKMAFPSGKVEGCMVQVTCGTTTLNGLWLDDVVYCPRHVICTSEDMLNPNYEDLLIR   60
P.2(Zeta)            SGFRKMAFPSGKVEGCMVQVTCGTTTLNGLWLDDVVYCPRHVICTSEDMLNPNYEDLLIR   60
B.1.1.529(Omicron)   SGFRKMAFPSGKVEGCMVQVTCGTTTLNGLWLDDVVYCPRHVICTSEDMLNPNYEDLLIR   60
                     **: **:.***.**.********: ********:.*:*****:* :::: :***: ***
                                                 90
SARS                 KSNHSFLVQA---GNVQLRVIGHSMQNCLLRLKVDTSNPKTPKYKFVRIQPGQTFSVLAC   117
MERS                 MTNHSFSVQKHIGAPANLRVVGHAMQGTLLKLTVDVANPSTPAYTFTTVKPGAAFSVLAC   120
WA1(COVID-19)        KSNHNFLVQA---GNVQLRVIGHSMQNCVLKLKVDTANPKTPKYKFVRIQPGQTFSVLAC   117
B.1.1.7(Alpha)       KSNHNFLVQA---GNVQLRVIGHSMQNCVLKLKVDTANPKTPKYKFVRIQPGQTFSVLAC   117
B.1.351(Beta)        KSNHNFLVQA---GNVQLRVIGHSMQNCVLKLRVDTANPKTPKYKFVRIQPGQTFSVLAC   117
B.1.427(Epsilon)     KSNHNFLVQA---GNVQLRVIGHSMQNCVLKLKVDTANPKTPKYKFVRIQPGQTFSVLAC   117
B.1.429(Epsilon)     KSNHNFLVQA---GNVQLRVIGHSMQNCVLKLKVDTANPKTPKYKFVRIQPGQTFSVLAC   117
B.1.525(Eta)         KSNHNFLVQA---GNVQLRVIGHSMQNCVLKLKVDTANPKTPKYKFVRIQPGQTFSVLAC   117
B.1.526(Iota)        KSNHNFLVQA---GNVQLRVIGHSMQNCVLKLKVDTANPKTPKYKFVRIQPGQTFSVLAC   117
B.1.526.1            KSNHNFLVQA---GNVQLRVIGHSMQNCVLKLKVDTANPKTPKYKFVRIQPGQTFSVLAC   117
B.1.617.1(Kappa)     KSNHNFLVQA---GNVQLRVIGHSMQNCVLKLKVDTANPKTPKYKFVRIQPGQTFSVLAC   117
B.1.617.2(Delta)     KSNHNFLVQA---GNVQLRVIGHSMQNCVLKLKVDTANPKTPKYKFVRIQPGQTFSVLAC   117
P.1(Gamma)           KSNHNFLVQA---GNVQLRVIGHSMQNCVLKLKVDTANPKTPKYKFVRIQPGQTFSVLAC   117
P.2(Zeta)            KSNHNFLVQA---GNVQLRVIGHSMQNCVLKLKVDTANPKTPKYKFVRIQPGQTFSVLAC   117
B.1.1.529(Omicron)   KSNHNFLVQA---GNVQLRVIGHSMQNCVLKLKVDTANPKTPKYKFVRIQPGQTFSVLAC   117
                      :**.* **    . .:***:**:**. :*:* **.:**.** *.*. ::** :******
                                  132
SARS                 YNGSPSGVYQCAMRPNHTIKGSFLNGSCGSVGFNIDYDCVSFCYMHHMELPTGVHAGTDL   177
MERS                 YNGRPTGTFTVVMRPNYTIKGSFLCGSCGSVGYTKEGSVINFCYMHQMELANGTHTGSAF   180
WA1(COVID-19)        YNGSPSGVYQCAMRPNFTIKGSFLNGSCGSVGFNIDYDCVSFCYMHHMELPTGVHAGTDL   177
B.1.1.7(Alpha)       YNGSPSGVYQCAMRPNFTIKGSFLNGSCGSVGFNIDYDCVSFCYMHHMELPTGVHAGTDL   177
B.1.351(Beta)        YNGSPSGVYQCAMRPNFTIKGSFLNGSCGSVGFNIDYDCVSFCYMHHMELPTGVHAGTDL   177
B.1.427(Epsilon)     YNGSPSGVYQCAMRPNFTIKGSFLNGSCGSVGFNIDYDCVSFCYMHHMELPTGVHAGTDL   177
B.1.429(Epsilon)     YNGSPSGVYQCAMRPNFTIKGSFLNGSCGSVGFNIDYDCVSFCYMHHMELPTGVHAGTDL   177
B.1.525(Eta)         YNGSPSGVYQCAMRPNFTIKGSFLNGSCGSVGFNIDYDCVSFCYMHHMELPTGVHAGTDL   177
B.1.526(Iota)        YNGSPSGVYQCAMRPNFTIKGSFLNGSCGSVGFNIDYDCVSFCYMHHMELPTGVHAGTDL   177
B.1.526.1            YNGSPSGVYQCAMRPNFTIKGSFLNGSCGSVGFNIDYDCVSFCYMHHMELPTGVHAGTDL   177
B.1.617.1(Kappa)     YNGSPSGVYQCAMRPNFTIKGSFLNGSCGSVGFNIDYDCVSFCYMHHMELPTGVHAGTDL   177
B.1.617.2(Delta)     YNGSPSGVYQCAMRPNFTIKGSFLNGSCGSVGFNIDYDCVSFCYMHHMELPTGVHAGTDL   177
P.1(Gamma)           YNGSPSGVYQCAMRPNFTIKGSFLNGSCGSVGFNIDYDCVSFCYMHHMELPTGVHAGTDL   177
P.2(Zeta)            YNGSPSGVYQCAMRPNFTIKGSFLNGSCGSVGFNIDYDCVSFCYMHHMELPTGVHAGTDL   177
B.1.1.529(Omicron)   YNGSPSGVYQCAMRHNFTIKGSFLNGSCGSVGFNIDYDCVSFCYMHHMELPTGVHAGTDL   177
                     *** *:*.:  .** *.******* *******:. : . :.*****:*** .*.*:*: :
                                             205
SARS                 EGKFYGPFVDRQTAQAAGTDTTITLNVLAWLYAAVINGDRWFLNRFTTTLNDFNLVAMKY   237
MERS                 DGTMYGAFMDKQVHQVQLTDKYCSVNVVAWLYAAILNGCAWFVKPNRTSVVSFNEWALAN   240
WA1(COVID-19)        EGNFYGPFVDRQTAQAAGTDTTITVNVLAWLYAAVINGDRWFLNRFTTTLNDFNLVAMKY   237
B.1.1.7(Alpha)       EGNFYGPFVDRQTAQAAGTDTTITVNVLAWLYAAVINGDRWFLNRFTTTLNDFNLVAMKY   237
B.1.351(Beta)        EGNFYGPFVDRQTAQAAGTDTTITVNVLAWLYAAVINGDRWFLNRFTTTLNDFNLVAMKY   237
B.1.427(Epsilon)     EGNFYGPFVDRQTAQAAGTDTTITVNVLAWLYAAVINGDRWFLNRFTTTLNDFNLVAMKY   237
B.1.429(Epsilon)     EGNFYGPFVDRQTAQAAGTDTTITVNVLAWLYAAVINGDRWFLNRFTTTLNDFNLVAMKY   237
B.1.525(Eta)         EGNFYGPFVDRQTAQAAGTDTTITVNVLAWLYAAVINGDRWFLNRFTTTLNDFNLVAMKY   237
B.1.526(Iota)        EGNFYGPFVDRQTAQAAGTDTTITVNVLAWLYAAVINGDRWFLNRFTTTLNDFNLVAMKY   237
B.1.526.1            EGNFYGPFVDRQTAQAAGTDTTITVNVLAWLYAAVINGDRWFLNRFTTTLNDFNLVAMKY   237
B.1.617.1(Kappa)     EGNFYGPFVDRQTAQAAGTDTTITVNVLAWLYAAVINGDRWFLNRFTTTLNDFNLVAMKY   237
B.1.617.2(Delta)     EGNFYGPFVDRQTAQAAGTDTTITVNVLAWLYAAVINGDRWFLNRFTTTLNDFNLVAMKY   237
P.1(Gamma)           EGNFYGPFVDRQTAQAAGTDTTITVNVLAWLYAAVINGDRWFLNRFTTTLNDFNLVAMKY   237
P.2(Zeta)            EGNFYGPFVDRQTAQAAGTDTTITVNVVAWLYAAVINGDRWFLNRFTTTLNDFNLVAMKY   237
B.1.1.529(Omicron)   EGNFYGPFVDRQTAQAAGTDTTITVNVLAWLYAAVINGDRWFLNRFTTTLNDFNLVAMKY   237
                     :*.:** *:*:*. *.  **.  ::**:******::**  **::   *:: .**  *:

SARS                 NYEPLTQDHVDILGPLSAQTGIAVLDMCAALKELLQNGMNGRTILGSTILEDEFTPFDVV   297
MERS                 QFTEFVGTQ--SVDMLAVKTGVAIEQLLYAI-QQLYTGFQGKQILGSTMLEDEFTPEDVN   297
WA1(COVID-19)        NYEPLTQDHVDILGPLSAQTGIAVLDMCASLKELLQNGMNGRTILGSALLEDEFTPFDVV   297
B.1.1.7(Alpha)       NYEPLTQDHVDILGPLSAQTGIAVLDMCASLKELLQNGMNGRTILGSALLEDEFTPFDVV   297
B.1.351(Beta)        NYEPLTQDHVDILGPLSAQTGIAVLDMCASLKELLQNGMNGRTILGSALLEDEFTPFDVV   297
B.1.427(Epsilon)     NYEPLTQDHVDILGPLSAQTGIAVLDMCASLKELLQNGMNGRTILGSALLEDEFTPFDVV   297
B.1.429(Epsilon)     NYEPLTQDHVDILGPLSAQTGIAVLDMCASLKELLQNGMNGRTILGSALLEDEFTPFDVV   297
B.1.525(Eta)         NYEPLTQDHVDILGPLSAQTGIAVLDMCASLKELLQNGMNGRTILGSALLEDEFTPFDVV   297
B.1.526(Iota)        NYEPLTQDHVDILGPLSAQTGIAVLDMCASLKELLQNGMNGRTILGSALLEDEFTPFDVV   297
B.1.526.1            NYEPLTQDHVDILGPLSAQTGIAVLDMCASLKELLQNGMNGRTILGSALLEDEFTPFDVV   297
B.1.617.1(Kappa)     NYEPLTQDHVDILGPLSAQTGIAVLDMCASLKELLQNGMNGRTILGSALLEDEFTPFDVV   297
B.1.617.2(Delta)     NYEPLTQDHVDILGPLSAQTGIAVLDMCASLKELLQNGMNGRTILGSALLEDEFTPFDVV   297
P.1(Gamma)           NYEPLTQDHVDILGPLSAQTGIAVLDMCASLKELLQNGMNGRTILGSALLEDEFTPFDVV   297
P.2(Zeta)            NYEPLTQDHVDILGPLSAQTGIAVLDMCASLKELLQNGMNGRTILGSALLEDEFTPFDVV   297
B.1.1.529(Omicron)   NYEPLTQDHVDILGPLSAQTGIAVLDMCASLKELLQNGMNGRTILGSALLEDEFTPFDVV   297
                     ::  :.  :   :. *:.:**:*: ::  :: : * .*::*: ****::******* **

SARS                 RQCSGVTFQ    306
MERS                 MQIMGVVMQ    306
WA1(COVID-19)        RQCSGVTFQ    306
B.1.1.7(Alpha)       RQCSGVTFQ    306
B.1.351(Beta)        RQCSGVTFQ    306
B.1.427(Epsilon)     RQCSGVTFQ    306
B.1.429(Epsilon)     RQCSGVTFQ    306
B.1.525(Eta)         RQCSGVTFQ    306
B.1.526(Iota)        RQCSGVTFQ    306
B.1.526.1            RQCSGVTFQ    306
B.1.617.1(Kappa)     RQCSGVTFQ    306
B.1.617.2(Delta)     RQCSGVTFQ    306
P.1(Gamma)           RQCSGVTFQ    306
P.2(Zeta)            RQCSGVTFQ    306
B.1.1.529(Omicron)   RQCSGVTFQ    306
                      *  **.:*
```

**Supplementary Figure S1**
