## Supplementary material for "Novel inhibitors against COVID-19 main protease suppressed viral infection": Figure S2

```
                          5        13            18-20      26
MERS               MIHSVFLLMFLLTPTESYVDVGPDSVKSACIEVDIQQTFFDKTWPRPIDVSKADGIIYPQ  60
SARS               ----MFIFLLFL-TLTSGSD------LDRCTTFDDVQ-----APNYTQHTSSMRGVYYPD  44
WA1(COVID-19)      ----MFVFLVLL-PLVSSQ----------CVNLTTRT-----QLPPAYTNSFTRGVYYPD  40
B.1.17(Alpha)      ----MFVFLVLL-PLVSSQ----------CVNLTTRT-----QLPPAYTNSFTRGVYYPD  40
B.1.351(Beta)      ----MFVFLVLL-PLVSSQ----------CVNFTTRT-----QLPPAYTNSFTRGVYYPD  40
B.1.427(Epsilon)   ----MFVFLVLL-PLVSIQ----------CVNLTTRT-----QLPPAYTNSFTRGVYYPD  40
B.1.429(Epsilon)   ----MFVFLVLL-PLVSIQ----------CVNLTTRT-----QLPPAYTNSFTRGVYYPD  40
B.1.525(Eta)       ----MFVFLVLL-PLVSSQ----------CVNLTTRT-----QLPPAYTNSFTRGVYYPD  40
B.1.526(Iota)      ----MFVFFVLL-PLVSSQ----------CVNLTTRT-----QLPPAYTNSFTRGVYYPD  40
B.1.526.1          ----MFVFLVLL-PLVSSQ----------CVNLTTRT-----QLPPAYTNSFTRGVYYPD  40
B.1.617.1(Kappa)   ----MFVFLVLL-PLVSSQ----------CVNLTTRT-----QLPPAYTNSFTRGVYYPD  40
B.1.617.2(Delta)   ----MFVFLVLL-PLVSSQ----------CVNLRTRT-----QLPPAYTNSFTRGVYYPD  40
P.1(Gamma)         ----MFVFLVLL-PLVSSQ----------CVNFTNRT-----QLPSAYTNSFTRGVYYPD  40
P.2(Zeta)          ----MFVFLVLL-PLVSSQ----------CVNLTTRT-----QLPPAYTNSFTRGVYYPD  40
B.1.1.529          ----MFVFLVLL-PLVSSQ----------CVNLTTRT-----QLPPAYTNSFTRGVYYPD  40
                       :*:::.:*     *            *  .                *   *: **:

                                52              67 69 70        80
MERS               GRTYSNITITYQGLF-PYQGDHGDMYVYSAGHATGTTPQKLFVANYSQDVKQFANGFVVR  119
SARS               EIFRSDTLYLTQDLFLPFYSNVTG---FHTIN-------HT----FGNPVIPFKDGIYFA  90
WA1(COVID-19)      KVFRSSVLHSTQDLFLPFFSNVTW---FHAIHVSGTNGTKR----FDNPVLPFNDGVYFA  93
B.1.17(Alpha)      KVFRSSVLHSTQDLFLPFFSNVTW---FHAI--SGTNGTKR----FDNPVLPFNDGVYFA  91
B.1.351(Beta)      KVFRSSVLHSTQDLFLPFFSNVTW---FHAIHVSGTNGTKR----FANPVLPFNDGVYFA  93
B.1.427(Epsilon)   KVFRSSVLHSTQDLFLPFFSNVTW---FHAIHVSGTNGTKR----FDNPVLPFNDGVYFA  93
B.1.429(Epsilon)   KVFRSSVLHSTQDLFLPFFSNVTW---FHAIHVSGTNGTKR----FDNPVLPFNDGVYFA  93
B.1.525(Eta)       KVFRSSVLHSTRDLFLPFFSNVTW---FHVI--SGTNGTKR----FDNPVLPFNDGVYFA  91
B.1.526(Iota)      KVFRSSVLHSTQDLFLPFFSNVTW---FHAIHVSGTNGTKR----FDNPVLPFNDGVYFA  93
B.1.526.1          KVFRSSVLHSTQDLFLPFFSNVTW---FHAIHVSGTNGTKR----FGNPVLPFNDGVYFA  93
B.1.617.1(Kappa)   KVFRSSVLHSTQDLFLPFFSNVTW---FHAIHVSGTNGTKR----FDNPVLPFNDGVYFA  93
B.1.617.2(Delta)   KVFRSSVLHSTQDLFLPFFSNVTW---FHAIHVSGTNGTTR----FDNPVLPFNDGVYFA  93
P.1(Gamma)         KVFRSSVLHSTQDLFLPFFSNVTW---FHAIHVSGTNGTKR----FDNPVLPFNDGVYFA  93
P.2(Zeta)          KVFRSSVLHSTQDLFLPFFSNVTW---FHAIHVSGTNGTKR----FDNPVLPFNDGVYFA  93
B.1.1.529          KVFRSSVLHSTQDLFLPFFSNVTW---FHVI--SGTNGTKR----FDNPVLPFNDGVYFA  91
                       *.     :.** *: .:        : .               : : *   * :*. .

                   95
MERS               IGAAANSTGTVIISPSTSATIRKIYPAFMLGSSVGNFSDGKMGRFFNHTLVLLPDGCGTL  179
SARS               AT-------------EK----SNVVRGWVFGSTMNNKSQ---------SVIIINNSTNVV  124
WA1(COVID-19)      ST-------------EK----SNIIRGWIFGTTLDSKTQ---------SLLIVNNATNVV  127
B.1.17(Alpha)      ST-------------EK----SNIIRGWIFGTTLDSKTQ---------SLLIVNNATNVV  125
B.1.351(Beta)      ST-------------EK----SNIIRGWIFGTTLDSKTQ---------SLLIVNNATNVV  127
B.1.427(Epsilon)   ST-------------EK----SNIIRGWIFGTTLDSKTQ---------SLLIVNNATNVV  127
B.1.429(Epsilon)   ST-------------EK----SNIIRGWIFGTTLDSKTQ---------SLLIVNNATNVV  127
B.1.525(Eta)       ST-------------EK----SNIIRGWIFGTTLDSKTQ---------SLLIVNNATNVV  125
B.1.526(Iota)      SI-------------EK----SNIIRGWIFGTTLDSKTQ---------SLLIVNNATNVV  127
B.1.526.1          ST-------------EK----SNIIRGWIFGTTLDSKTQ---------SLLIVNNATNVV  127
B.1.617.1(Kappa)   ST-------------EK----SNIIRGWIFGTTLDSKTQ---------SLLIVNNATNVV  127
B.1.617.2(Delta)   ST-------------EK----SNIIRGWIFGTTLDSKTQ---------SLLIVNNATNVV  127
P.1(Gamma)         ST-------------EK----SNIIRGWIFGTTLDSKTQ---------SLLIVNNATNVV  127
P.2(Zeta)          ST-------------EK----SNIIRGWIFGTTLDSKTQ---------SLLIVNNATNVV  127
B.1.1.529          SI-------------EK----SNIIRGWIFGTTLDSKTQ---------SLLIVNNATNVV  125
                                  ..    ::  .:::*:::.. ::         ::::: :.  ..:

                                              138  142-145    152 154  156-158
MERS               LRAFYCILEPRSGNHCPAGNSYTSFATYHTPATDCSDGNYNRNASLNSFKEYFNLRNCTF  239
SARS               IRA--CNFE-----LC------------DNPFFAVSKPMGT----QTHTMIFDNAFNCTF  161
WA1(COVID-19)      IKV--CEFQ-----FC------------NDPFLGVYYHKNNKSWMESEFRVYSSANNCTF  168
B.1.17(Alpha)      IKV--CEFQ-----FC------------NDPFLGV-YHKNNKSWMESEFRVYSSANNCTF  165
B.1.351(Beta)      IKV--CEFQ-----FC------------NDPFLGVYYHKNNKSWMESEFRVYSSANNCTF  168
B.1.427(Epsilon)   IKV--CEFQ-----FC------------NDPFLGVYHHKNNKSCMESEFRVYSSANNCTF  168
B.1.429(Epsilon)   IKV--CEFQ-----FC------------NDPFLGVYYHKNNKSCMESEFRVYSSANNCTF  168
B.1.525(Eta)       IKV--CEFQ-----FC------------NDPFLGVY-HKNNKSWMESEFRVYSSANNCTF  165
B.1.526(Iota)      IKV--CEFQ-----FC------------NDPFLGVYYHKNNKSWMESEFRVYSSANNCTF  168
B.1.526.1          IKV--CEFQ-----FC------------NDPFLGVY-HKNNKSWMESESRVYSSANNCTF  167
B.1.617.1(Kappa)   IKV--CEFQ-----FC------------NDPFLGVYYHKNNKSWMKSEFRVYSSANNCTF  168
B.1.617.2(Delta)   IKV--CEFQ-----FC------------NDPFLGVYYHKNNKSWMES--GVYSSANNCTF  166
P.1(Gamma)         IKV--CEFQ-----FC------------NYPFLGVYYHKNNKSWMESEFRVYSSANNCTF  168
P.2(Zeta)          IKV--CEFQ-----FC------------NDPFLGVYYHKNNKSWMESEFRVYSSANNCTF  168
B.1.1.529          IKV--CEFQ-----FC------------NDPFLD---HKNNKSWMESEFRVYSSANNCTF  163
                   ::.  * ::     *             . *          .     .    : .  ****
```

```
                                          190                   211 - 217
MERS                MYTYNITEDEIL------------------EWFGITQTAQGVHLFSSRYVDLYGG--NMF   279
SARS                EYISDAFSLDVSEKSGNFKHLREFVFKNKDGFLYVYKGYQPIDV--V--RDLPSGFNTLK   217
WA1(COVID-19)       EYVSQPFLMDLEGKQGNFKNLREFVFKNIDGYFKIYSKHTPINL--V--RDLPQGFSALE   224
B.1.17(Alpha)       EYVSQPFLMDLEGKQGNFKNLREFVFKNIDGYFKIYSKHTPINL--V--RDLPQGFSALE   221
B.1.351(Beta)       EYVSQPFLMDLEGKQGNFKNLREFVFKNIDGYFKIYSKHTPINL--V--RGLPQGFSALE   224
B.1.427(Epsilon)    EYVSQPFLMDLEGKQGNFKNLREFVFKNIDGYFKIYSKHTPINL--V--RDLPQGFSALE   224
B.1.429(Epsilon)    EYVSQPFLMDLEGKQGNFKNLREFVFKNIDGYFKIYSKHTPINL--V--RDLPQGFSALE   224
B.1.525(Eta)        EYVSQPFLMDLEGKQGNFKNLREFVFKNIDGYFKIYSKHTPINL--V--RDLPQGFSALE   221
B.1.526(Iota)       EYVSQPFLMDLEGKQGNFKNLREFVFKNIDGYFKIYSKHTPINL--V--RDLPQGFSALE   224
B.1.526.1           EYVSQPFLMDLEGKQGNFKNLREFVFKNIDGYFKIYSKHTPINL--V--RDLPQGFSALE   223
B.1.617.1(Kappa)    EYVSQPFLMDLEGKQGNFKNLREFVFKNIDGYFKIYSKHTPINL--V--RDLPQGFSALE   224
B.1.617.2(Delta)    EYVSQPFLMDLEGKQGNFKNLREFVFKNIDGYFKIYSKHTPINL--V--RDLPQGFSALE   222
P.1(Gamma)          EYVSQPFLMDLEGKQGNFKNLSEFVFKNIDGYFKIYSKHTPINL--V--RDLPQGFSALE   224
P.2(Zeta)           EYVSQPFLMDLEGKQGNFKNLREFVFKNIDGYFKIYSKHTPINL--V--RDLPQGFSALE   224
B.1.1.529           EYVSQPFLMDLEGKQGNFKNLREFVFKNIDGYFKIYSKHTPIIVREP--EDLPQGFSALE   221
                     *  :    ::                    :: : .    : :      .*  *   :

                                     241-243
MERS                QFATLPVYDTIKYYSIIPHSIR---SIQSDRKAW----AAFYVYKLQPLTFLLDFSVDGY   332
SARS                PIFKLPLGINITNFRAILTAFS------PAQDIWGTSAAAYFVGYLKPTTFMLKYDENGT   271
WA1(COVID-19)       PLVDLPIGINITRFQTLLALHRSYLTPGDSSSGWTAGAAAYYVGYLQPRTFLLKYNENGT   284
B.1.17(Alpha)       PLVDLPIGINITRFQTLLALHRSYLTPGDSSSGWTAGAAAYYVGYLQPRTFLLKYNENGT   281
B.1.351(Beta)       PLVDLPIGINITRFQT---LHRSYLTPGDSSSGWTAGAAAYYVGYLQPRTFLLKYNENGT   281
B.1.427(Epsilon)    PLVDLPIGINITRFQTLLALHRSYLTPGDSSSGWTAGAAAYYVGYLQPRTFLLKYNENGT   284
B.1.429(Epsilon)    PLVDLPIGINITRFQTLLALHRSYLTPGDSSSGWTAGAAAYYVGYLQPRTFLLKYNENGT   284
B.1.525(Eta)        PLVDLPIGINITRFQTLLALHRSYLTPGDSSSGWTAGAAAYYVGYLQPRTFLLKYNENGT   281
B.1.526(Iota)       PLVDLPIGINITRFQTLLALHRSYLTPGGSSSGWTAGAAAYYVGYLQPRTFLLKYNENGT   284
B.1.526.1           PLVDLPIGINITRFQTLLALHRSYLTPGDSSSGWTAGAAAYYVGYLQPRTFLLKYNENGT   283
B.1.617.1(Kappa)    PLVDLPIGINITRFQTLLALHRSYLTPGDSSSGWTAGAAAYYVGYLQPRTFLLKYNENGT   284
B.1.617.2(Delta)    PLVDLPIGINITRFQTLLALHRSYLTPGDSSSGWTAGAAAYYVGYLQPRTFLLKYNENGT   282
P.1(Gamma)          PLVDLPIGINITRFQTLLALHRSYLTPGDSSSGWTAGAAAYYVGYLQPRTFLLKYNENGT   284
P.2(Zeta)           PLVDLPIGINITRFQTLLALHRSYLTPGDSSSGWTAGAAAYYVGYLQPRTFLLKYNENGT   284
B.1.1.529           PLVDLPIGINITRFQTLLALHRSYLTPGDSSSGWTAGAAAYYVGYLQPRTFLLKYNENGT   281
                     :  **:  .*. :                 . *    **::*  *:* **:*.:. :*

                                                                  339
MERS                IRRAIDCGFNDLSQLHCSYESFDVESGVYSVSSFEAKPSGSVVEQAEG-VECDFSPLLSG   391
SARS                ITDAVDCSQNPLAELKCSVKSFEIDKGIYQTSNFRVVPSGDVVRFPNITNLCPFGEVFNA   331
WA1(COVID-19)       ITDAVDCALDPLSETKCTLKSFTVEKGIYQTSNFRVQPTESIVRFPNITNLCPFGEVFNA   344
B.1.17(Alpha)       ITDAVDCALDPLSETKCTLKSFTVEKGIYQTSNFRVQPTESIVRFPNITNLCPFGEVFNA   341
B.1.351(Beta)       ITDAVDCALDPLSETKCTLKSFTVEKGIYQTSNFRVQPTESIVRFPNITNLCPFGEVFNA   341
B.1.427(Epsilon)    ITDAVDCALDPLSETKCTLKSFTVEKGIYQTSNFRVQPTESIVRFPNITNLCPFGEVFNA   344
B.1.429(Epsilon)    ITDAVDCALDPLSETKCTLKSFTVEKGIYQTSNFRVQPTESIVRFPNITNLCPFGEVFNA   344
B.1.525(Eta)        ITDAVDCALDPLSETKCTLKSFTVEKGIYQTSNFRVQPTESIVRFPNITNLCPFGEVFNA   341
B.1.526(Iota)       ITDAVDCALDPLSETKCTLKSFTVEKGIYQTSNFRVQPTESIVRFPNITNLCPFGEVFNA   344
B.1.526.1           ITDAVDCALDPLSETKCTLKSFTVEKGIYQTSNFRVQPTESIVRFPNITNLCPFGEVFNA   343
B.1.617.1(Kappa)    ITDAVDCALDPLSETKCTLKSFTVEKGIYQTSNFRVQPTESIVRFPNITNLCPFGEVFNA   344
B.1.617.2(Delta)    ITDAVDCALDPLSETKCTLKSFTVEKGIYQTSNFRVQPTESIVRFPNITNLCPFGEVFNA   342
P.1(Gamma)          ITDAVDCALDPLSETKCTLKSFTVEKGIYQTSNFRVQPTESIVRFPNITNLCPFGEVFNA   344
P.2(Zeta)           ITDAVDCALDPLSETKCTLKSFTVEKGIYQTSNFRVQPTESIVRFPNITNLCPFGEVFNA   344
B.1.1.529           ITDAVDCALDPLSETKCTLKSFTVEKGIYQTSNFRVQPTESIVRFPNITNLCPFDEVFNA   341
                    *  *:**. : *:: :*: :** ::.*:*..*.*.. *: .:*.  :    * *. ::..

                                         371 373 375
MERS                -TPPQVYNFKRLVFTNCNYNLTKLLSLFSVNDFTCSQISPAAIASNCYSSLILDYFSYPL   450
SARS                TKFPSVYAWERKKISNCVADYSVLYNSTFFSTFKCYGVSATKLNDLCFSNVYADSFVVKG   391
WA1(COVID-19)       TRFASVYAWNRKRISNCVADYSVLYNSASFSTFKCYGVSPTKLNDLCFTNVYADSFVIRG   404
B.1.17(Alpha)       TRFASVYAWNRKRISNCVADYSVLYNSASFSTFKCYGVSPTKLNDLCFTNVYADSFVIRG   401
B.1.351(Beta)       TRFASVYAWNRKRISNCVADYSVLYNSASFSTFKCYGVSPTKLNDLCFTNVYADSFVIRG   401
B.1.427(Epsilon)    TRFASVYAWNRKRISNCVADYSVLYNSASFSTFKCYGVSPTKLNDLCFTNVYADSFVIRG   404
B.1.429(Epsilon)    TRFASVYAWNRKRISNCVADYSVLYNSASFSTFKCYGVSPTKLNDLCFTNVYADSFVIRG   404
B.1.525(Eta)        TRFASVYAWNRKRISNCVADYSVLYNSASFSTFKCYGVSPTKLNDLCFTNVYADSFVIRG   401
B.1.526(Iota)       TRFASVYAWNRKRISNCVADYSVLYNSASFSTFKCYGVSPTKLNDLCFTNVYADSFVIRG   404
B.1.526.1           TRFASVYAWNRKRISNCVADYSVLYNSASFSTFKCYGVSPTKLNDLCFTNVYADSFVIRG   403
B.1.617.1(Kappa)    TRFASVYAWNRKRISNCVADYSVLYNSASFSTFKCYGVSPTKLNDLCFTNVYADSFVIRG   404
B.1.617.2(Delta)    TRFASVYAWNRKRISNCVADYSVLYNSASFSTFKCYGVSPTKLNDLCFTNVYADSFVIRG   402
P.1(Gamma)          TRFASVYAWNRKRISNCVADYSVLYNSASFSTFKCYGVSPTKLNDLCFTNVYADSFVIRG   404
P.2(Zeta)           TRFASVYAWNRKRISNCVADYSVLYNSASFSTFKCYGVSPTKLNDLCFTNVYADSFVIRG   404
B.1.1.529           TRFASVYAWNRKRISNCVADYSVLYNLAPFFTFKCYGVSPTKLNDLCFTNVYADSFVIRG   401
                        .** ::*  ::**  : : * .   .  *.*  :* : : . *::.:  * *
```

```
                            417                            440    446     452
MERS              SMKSDLSVSSAGPISQFNYKQSFSNPTCLILATVPHNLTTITKPLKYSYINKCSRLLSDD   510
SARS              DDVRQIAPGQTGVIADYNYKLPDDFMGCVLAWNTRNIDATSTGNYNYKYRYLRHGKLRPF   451
WA1(COVID-19)     DEVRQIAPGQTGKIADYNYKLPDDFTGCVIAWNSNNLDSKVGGNYNYLYRLFRKSNLKPF   464
B.1.17(Alpha)     DEVRQIAPGQTGKIADYNYKLPDDFTGCVIAWNSNNLDSKVGGNYNYLYRLFRKSNLKPF   461
B.1.351(Beta)     DEVRQIAPGQTGNIADYNYKLPDDFTGCVIAWNSNNLDSKVGGNYNYLYRLFRKSNLKPF   461
B.1.427(Epsilon)  DEVRQIAPGQTGKIADYNYKLPDDFTGCVIAWNSNNLDSKVGGNYNYRYRLFRKSNLKPF   464
B.1.429(Epsilon)  DEVRQIAPGQTGKIADYNYKLPDDFTGCVIAWNSNNLDSKVGGNYNYRYRLFRKSNLKPF   464
B.1.525(Eta)      DEVRQIAPGQTGKIADYNYKLPDDFTGCVIAWNSNNLDSKVGGNYNYLYRLFRKSNLKPF   461
B.1.526(Iota)     DEVRQIAPGQTGKIADYNYKLPDDFTGCVIAWNSNNLDSKVGGNYNYLYRLFRKSNLKPF   464
B.1.526.1         DEVRQIAPGQTGKIADYNYKLPDDFTGCVIAWNSNNLDSKVGGNYNYRYRLFRKSNLKPF   463
B.1.617.1(Kappa)  DEVRQIAPGQTGKIADYNYKLPDDFTGCVIAWNSNNLDSKVGGNYNYRYRLFRKSNLKPF   464
B.1.617.2(Delta)  DEVRQIAPGQTGKIADYNYKLPDDFTGCVIAWNSNNLDSKVGGNYNYRYRLFRKSNLKPF   462
P.1(Gamma)        DEVRQIAPGQTGTIADYNYKLPDDFTGCVIAWNSNNLDSKVGGNYNYLYRLFRKSNLKPF   464
P.2(Zeta)         DEVRQIAPGQTGKIADYNYKLPDDFTGCVIAWNSNNLDSKVGGNYNYLYRLFRKSNLKPF   464
B.1.1.529         DEVRQIAPGQTGNIADYNYKLPDDFTGCVIAWNSNKLDSKVSGNYNYLYRLFRKSNLKPF   461
                  .   ::: ..:* *:::***   .   *::  .  :  :.      :* *       *

                           477 478  484                493 496 498 501 505
MERS              RTEVPQLVNANQYSPCVSIVPSTVWEDGDYYRKQLSPLEGGGWLVASGSTVAMTEQLQMG   570
SARS              ERDISNVPFSPDGKPCTP-PALNCY----------WPLNDYGFYTTTGIGYQPYRVVVLS   500
WA1(COVID-19)     ERDISTEIYQAGSTPCNGVEGFNCY----------FPLQSYGFQPTNGVGYQPYRVVVLS   514
B.1.17(Alpha)     ERDISTEIYQAGSTPCNGVEGFNCY----------FPLQSYGFQPTYGVGYQPYRVVVLS   511
B.1.351(Beta)     ERDISTEIYQAGSTPCNGVKGFNCY----------FPLQSYGFQPTYGVGYQPYRVVVLS   511
B.1.427(Epsilon)  ERDISTEIYQAGSTPCNGVEGFNCY----------FPLQSYGFQPTNGVGYQPYRVVVLS   514
B.1.429(Epsilon)  ERDISTEIYQAGSTPCNGVEGFNCY----------FPLQSYGFQPTNGVGYQPYRVVVLS   514
B.1.525(Eta)      ERDISTEIYQAGSTPCNGVKGFNCY----------FPLQSYGFQPTNGVGYQPYRVVVLS   511
B.1.526(Iota)     ERDISTEIYQAGSTPCNGVKGFNCY----------FPLQSYGFQPTNGVGYQPYRVVVLS   514
B.1.526.1         ERDISTEIYQAGSTPCNGVEGFNCY----------FPLQSYGFQPTNGVGYQPYRVVVLS   513
B.1.617.1(Kappa)  ERDISTEIYQAGSTPCNGVQGFNCY----------FPLQSYGFQPTNGVGYQPYRVVVLS   514
B.1.617.2(Delta)  ERDISTEIYQAGSKPCNGVEGFNCY----------FPLQSYGFQPTNGVGYQPYRVVVLS   512
P.1(Gamma)        ERDISTEIYQAGSTPCNGVKGFNCY----------FPLQSYGFQPTYGVGYQPYRVVVLS   514
P.2(Zeta)         ERDISTEIYQAGSTPCNGVKGFNCY----------FPLQSYGFQPTNGVGYQPYRVVVLS   514
B.1.1.529         ERDISTEIYQAGNKPCNGVAGFNCY----------FPLRSYSFRPTYGVGHQPYRVVVLS   511
                  . ::         .**     . :            **.. .: : *     . : :.
```

```
                                                           547
MERS              FGITVQYGTDTNSVCPKLEFANDTKIASQLGNCVEYSLYGVSGRGVFQNCTAVGVRQQRF   630
SARS              FEL----LNAPATVCGP-----KLSTDLIKNQCVNFNFNGLTGTGVLTPSSKRFQPFQQF   551
WA1(COVID-19)     FEL----LHAPATVCGP-----KKSTNLVKNKCVNFNFNGLTGTGVLTESNKKFLPFQQF   565
B.1.17(Alpha)     FEL----LHAPATVCGP-----KKSTNLVKNKCVNFNFNGLTGTGVLTESNKKFLPFQQF   562
B.1.351(Beta)     FEL----LHAPATVCGP-----KKSTNLVKNKCVNFNFNGLTGTGVLTESNKKFLPFQQF   562
B.1.427(Epsilon)  FEL----LHAPATVCGP-----KKSTNLVKNKCVNFNFNGLTGTGVLTESNKKFLPFQQF   565
B.1.429(Epsilon)  FEL----LHAPATVCGP-----KKSTNLVKNKCVNFNFNGLTGTGVLTESNKKFLPFQQF   565
B.1.525(Eta)      FEL----LHAPATVCGP-----KKSTNLVKNKCVNFNFNGLTGTGVLTESNKKFLPFQQF   562
B.1.526(Iota)     FEL----LHAPATVCGP-----KKSTNLVKNKCVNFNFNGLTGTGVLTESNKKFLPFQQF   565
B.1.526.1         FEL----LHAPATVCGP-----KKSTNLVKNKCVNFNFNGLTGTGVLTESNKKFLPFQQF   564
B.1.617.1(Kappa)  FEL----LHAPATVCGP-----KKSTNLVKNKCVNFNFNGLTGTGVLTESNKKFLPFQQF   565
B.1.617.2(Delta)  FEL----LHAPATVCGP-----KKSTNLVKNKCVNFNFNGLTGTGVLTESNKKFLPFQQF   563
P.1(Gamma)        FEL----LHAPATVCGP-----KKSTNLVKNKCVNFNFNGLTGTGVLTESNKKFLPFQQF   565
P.2(Zeta)         FEL----LHAPATVCGP-----KKSTNLVKNKCVNFNFNGLTGTGVLTESNKKFLPFQQF   565
B.1.1.529         FEL----LHAPATVCGP-----KKSTNLVKNKCVNFNFNGLKGTGVLTESNKKFLPFQQF   562
                  * :          :**        . .     .:**::.: *:.* **:   ..       *:*

                     570                                                614
MERS              VYDAYQNLVGYYSD--DGNYYCLRACVSVPVSVIYD--KETKTHATLFGSVACEHISSTM   686
SARS              GRDVSDFTD-SVRDPKTSEILDISPCSFGGVSVITPGTNASSEVAVLYQDVNCTDVSTAI   610
WA1(COVID-19)     GRDIADTTD-AVRDPQTLEILDITPCSFGGVSVITPGTNTSNQVAVLYQDVNCTEVPVAI   624
B.1.17(Alpha)     GRDIDDTTD-AVRDPQTLEILDITPCSFGGVSVITPGTNTSNQVAVLYQGVNCTEVPVAI   621
B.1.351(Beta)     GRDIADTTD-AVRDPQTLEILDITPCSFGGVSVITPGTNTSNQVAVLYQGVNCTEVPVAI   621
B.1.427(Epsilon)  GRDIADTTD-AVRDPQTLEILDITPCSFGGVSVITPGTNTSNQVAVLYQGVNCTEVPVAI   624
B.1.429(Epsilon)  GRDIADTTD-AVRDPQTLEILDITPCSFGGVSVITPGTNTSNQVAVLYQGVNCTEVPVAI   624
B.1.525(Eta)      GRDIADTTD-AVRDPQTLEILDITPCSFGGVSVITPGTNTSNQVAVLYQGVNCTEVPVAI   621
B.1.526(Iota)     GRDIADTTD-AVRDPQTLEILDITPCSFGGVSVITPGTNTSNQVAVLYQGVNCTEVPVAI   624
B.1.526.1         GRDIADTTD-AVRDPQTLEILDITPCSFGGVSVITPGTNTSNQVAVLYQGVNCTEVPVAI   623
B.1.617.1(Kappa)  GRDIADTTD-AVRDPQTLEILDITPCSFGGVSVITPGTNTSNQVAVLYQGVNCTEVPVAI   624
B.1.617.2(Delta)  GRDIADTTD-AVRDPQTLEILDITPCSFGGVSVITPGTNTSNQVAVLYQGVNCTEVPVAI   622
P.1(Gamma)        GRDIADTTD-AVRDPQTLEILDITPCSFGGVSVITPGTNTSNQVAVLYQGVNCTEVPVAI   624
P.2(Zeta)         GRDIADTTD-AVRDPQTLEILDITPCSFGGVSVITPGTNTSNQVAVLYQGVNCTEVPVAI   624
B.1.1.529         GRDIADTTD-AVRDPQTLEILDITPCSFGGVSVITPGTNTSNQVAVLYQGVNCTEVPVAI   621
                   *   :        *    :    :   *     ****     : :.    *.*: .* *  .:   ::
```

```
                                                   655                  677 679
MERS                SQYSRSTRSMLKRRDSTYGPLQTPVGCVLGLVNSSLFVEDCKLPLGQSLCALPDTPSTLT   746
SARS                HADQLT--PAWRIYSTGNNVFQTQAGCLIGAEHVD-TSYECDIPIGAGICASYHTVSL--   665
WA1(COVID-19)       HADQLT--PTWRVYSTGSNVFQTRAGCLIGAEHVN-NSYECDIPIGAGICASYQTQTN-S   680
B.1.17(Alpha)       HADQLT--PTWRVYSTGSNVFQTRAGCLIGAEHVN-NSYECDIPIGAGICASYQTQTN-S   677
B.1.351(Beta)       HADQLT--PTWRVYSTGSNVFQTRAGCLIGAEHVN-NSYECDIPIGAGICASYQTQTN-S   677
B.1.427(Epsilon)    HADQLT--PTWRVYSTGSNVFQTRAGCLIGAEHVN-NSYECDIPIGAGICASYQTQTN-S   680
B.1.429(Epsilon)    HADQLT--PTWRVYSTGSNVFQTRAGCLIGAEHVN-NSYECDIPIGAGICASYQTQTN-S   680
B.1.525(Eta)        HADQLT--PTWRVYSTGSNVFQTRAGCLIGAEHVN-NSYECDIPIGAGICASYQTHTN-S   677
B.1.526(Iota)       HADQLT--PTWRVYSTGSNVFQTRAGCLIGAEHVN-NSYECDIPIGAGICASYQTQTN-S   680
B.1.526.1           HADQLT--PTWRVYSTGSNVFQTRAGCLIGAEHVN-NSYECDIPIGAGICASYQTQTN-S   679
B.1.617.1(Kappa)    HADQLT--PTWRVYSTGSNVFQTRAGCLIGAEHVN-NSYECDIPIGAGICASYQTQTN-S   680
B.1.617.2(Delta)    HADQLT--PTWRVYSTGSNVFQTRAGCLIGAEHVN-NSYECDIPIGAGICASYQTQTN-S   678
P.1(Gamma)          HADQLT--PTWRVYSTGSNVFQTRAGCLIGAEYVN-NSYECDIPIGAGICASYQTQTN-S   680
P.2(Zeta)           HADQLT--PTWRVYSTGSNVFQTRAGCLIGAEHVN-NSYECDIPIGAGICASYQTQTN-S   680
B.1.1.529           HADQLT--PTWRVYSTGSNVFQTRAGCLIGAEYVN-NSYECDIPIGAGICASYQTQTK-S   677
                       . :     :  .:  . :** .**::*    .    :*.:*:* .:**  .* :

                    681                   701            716
MERS                PRSVRSVPGEMRLASIAFNHPIQV-DQLNSSYFKLSIPTNFSFGVTQEYIQTTIQKVTVD   805
SARS                ---LRSTSQKSI---VAYTMSLGADSSIAYSNNTIAIPTNFSISITTEVMPVSMAKTSVD   719
WA1(COVID-19)       PRRARSVASQSI---IAYTMSLGAENSVAYSNNSIAIPTNFTISVTTEILPVSMTKTSVD   737
B.1.17(Alpha)       HRRARSVASQSI---IAYTMSLGAENSVAYSNNSIAIPINFTISVTTEILPVSMTKTSVD   734
B.1.351(Beta)       PRRARSVASQSI---IAYTMSLGVENSVAYSNNSIAIPTNFTISVTTEILPVSMTKTSVD   734
B.1.427(Epsilon)    PRRARSVASQSI---IAYTMSLGAENSVAYSNNSIAIPTNFTISVTTEILPVSMTKTSVD   737
B.1.429(Epsilon)    PRRARSVASQSI---IAYTMSLGAENSVAYSNNSIAIPTNFTISVTTEILPVSMTKTSVD   737
B.1.525(Eta)        PRRARSVASQSI---IAYTMSLGAENSVAYSNNSIAIPTNFTISVTTEILPVSMTKTSVD   734
B.1.526(Iota)       PRRARSVASQSI---IAYTMSLGVENSVAYSNNSIAIPTNFTISVTTEILPVSMTKTSVD   737
B.1.526.1           PRRARSVASQSI---IAYTMSLGAENSVAYSNNSIAIPTNFTISVTTEILPVSMTKTSVD   736
B.1.617.1(Kappa)    RRRARSVASQSI---IAYTMSLGAENSVAYSNNSIAIPTNFTISVTTEILPVSMTKTSVD   737
B.1.617.2(Delta)    RRRARSVASQSI---IAYTMSLGAENSVAYSNNSIAIPTNFTISVTTEILPVSMTKTSVD   735
P.1(Gamma)          PRRARSVASQSI---IAYTMSLGAENSVAYSNNSIAIPTNFTISVTTEILPVSMTKTSVD   737
P.2(Zeta)           PRRARSVASQSI---IAYTMSLGAENSVAYSNNSIAIPTNFTISVTTEILPVSMTKTSVD   737
B.1.1.529           HRRARSVASQSI---IAYTMSLGAENSVAYSNNSIAIPTNFTISVTTEILPVSMTKTSVD   734
                        **.  :     :*:.  : . ..:  *  .::** **::.:* * : .:: *.:**

                                             764                            796
MERS                CKQYVCNGFQKCEQLLREYGQFCSKINQALHGANLRQDDSVRNLFASVKSSQSSPIIPGF   865
SARS                CNMYICGDSTECANLLLQYGSFCTQLNRALSGIAAEQDRNTREVFAQVKQMYKTPTLKYF   779
WA1(COVID-19)       CTMYICGDSTECSNLLLQYGSFCTQLNRALTGIAVEQDKNTQEVFAQVKQIYKTPPIKDF   797
B.1.17(Alpha)       CTMYICGDSTECSNLLLQYGSFCTQLNRALTGIAVEQDKNTQEVFAQVKQIYKTPPIKDF   794
B.1.351(Beta)       CTMYICGDSTECSNLLLQYGSFCTQLNRALTGIAVEQDKNTQEVFAQVKQIYKTPPIKDF   794
B.1.427(Epsilon)    CTMYICGDSTECSNLLLQYGSFCTQLNRALTGIAVEQDKNTQEVFAQVKQIYKTPPIKDF   797
B.1.429(Epsilon)    CTMYICGDSTECSNLLLQYGSFCTQLNRALTGIAVEQDKNTQEVFAQVKQIYKTPPIKDF   797
B.1.525(Eta)        CTMYICGDSTECSNLLLQYGSFCTQLNRALTGIAVEQDKNTQEVFAQVKQIYKTPPIKDF   794
B.1.526(Iota)       CTMYICGDSTECSNLLLQYGSFCTQLNRALTGIAVEQDKNTQEVFAQVKQIYKTPPIKDF   797
B.1.526.1           CTMYICGDSTECSNLLLQYGSFCTQLNRALTGIAVEQDKNTQEVFAQVKQIYKTPPIKDF   796
B.1.617.1(Kappa)    CTMYICGDSTECSNLLLQYGSFCTQLNRALTGIAVEQDKNTQEVFAQVKQIYKTPPIKDF   797
B.1.617.2(Delta)    CTMYICGDSTECSNLLLQYGSFCTQLNRALTGIAVEQDKNTQEVFAQVKQIYKTPPIKDF   795
P.1(Gamma)          CTMYICGDSTECSNLLLQYGSFCTQLNRALTGIAVEQDKNTQEVFAQVKQIYKTPPIKDF   797
P.2(Zeta)           CTMYICGDSTECSNLLLQYGSFCTQLNRALTGIAVEQDKNTQEVFAQVKQIYKTPPIKDF   797
B.1.1.529           CTMYICGDSTECSNLLLQYGSFCTQLKRALTGIAVEQDKNTQEVFAQVKQIYKTPPIKYF   794
                    *. *:*..  :* :** :**.**:::::** *   .** ..:::**.**.  .:* :  *

MERS                GGDFNLTLLEPVSISTGSRSARSAIEDLLFDKVTIADPGYMQGYDDCMQQGPASARDLIC   925
SARS                GGF-NFSQILPD---PLKPTKRSFIEDLLFNKVTLADAGFMKQYGECL--GDINARDLIC   833
WA1(COVID-19)       GGF-NFSQILPD---PSKPSKRSFIEDLLFNKVTLADAGFIKQYGDCL--GDIAARDLIC   851
B.1.17(Alpha)       GGF-NFSQILPD---PSKPSKRSFIEDLLFNKVTLADAGFIKQYGDCL--GDIAARDLIC   848
B.1.351(Beta)       GGF-NFSQILPD---PSKPSKRSFIEDLLFNKVTLADAGFIKQYGDCL--GDIAARDLIC   848
B.1.427(Epsilon)    GGF-NFSQILPD---PSKPSKRSFIEDLLFNKVTLADAGFIKQYGDCL--GDIAARDLIC   851
B.1.429(Epsilon)    GGF-NFSQILPD---PSKPSKRSFIEDLLFNKVTLADAGFIKQYGDCL--GDIAARDLIC   851
B.1.525(Eta)        GGF-NFSQILPD---PSKPSKRSFIEDLLFNKVTLADAGFIKQYGDCL--GDIAARDLIC   848
B.1.526(Iota)       GGF-NFSQILPD---PSKPSKRSFIEDLLFNKVTLADAGFIKQYGDCL--GDIAARDLIC   851
B.1.526.1           GGF-NFSQILPD---PSKPSKRSFIEDLLFNKVTLADAGFIKQYGDCL--GDIAARDLIC   850
B.1.617.1(Kappa)    GGF-NFSQILPD---PSKPSKRSFIEDLLFNKVTLADAGFIKQYGDCL--GDIAARDLIC   851
B.1.617.2(Delta)    GGF-NFSQILPD---PSKPSKRSFIEDLLFNKVTLADAGFIKQYGDCL--GDIAARDLIC   849
P.1(Gamma)          GGF-NFSQILPD---PSKPSKRSFIEDLLFNKVTLADAGFIKQYGDCL--GDIAARDLIC   851
P.2(Zeta)           GGF-NFSQILPD---PSKPSKRSFIEDLLFNKVTLADAGFIKQYGDCL--GDIAARDLIC   851
B.1.1.529           GGF-NFSQILPD---PSKPSKRSFIEDLLFNKVTLADAGFIKQYGDCL--GDIAARDLIC   848
                    **  *:: : *      . : ** ******:***:** *::: *.:*:  *   ******
```

```
                              856                           888
MERS                 AQYVAGYKVLPPLMDVNMEAAYTSSLLGSIAGVGWTAGLSSFAAIPFAQSIFYRLNGVGI    985
SARS                 AQKFNGLTVLPPLLTDDMIAAYTAALVSGTATAGWTFGAGAALQIPFAMQMAYRFNGIGV    893
WA1(COVID-19)        AQKFNGLTVLPPLLTDEMIAQYTSALLAGTITSGWTFGAGAALQIPFAMQMAYRFNGIGV    911
B.1.17(Alpha)        AQKFNGLTVLPPLLTDEMIAQYTSALLAGTITSGWTFGAGAALQIPFAMQMAYRFNGIGV    908
B.1.351(Beta)        AQKFNGLTVLPPLLTDEMIAQYTSALLAGTITSGWTFGAGAALQIPFAMQMAYRFNGIGV    908
B.1.427(Epsilon)     AQKFNGLTVLPPLLTDEMIAQYTSALLAGTITSGWTFGAGAALQIPFAMQMAYRFNGIGV    911
B.1.429(Epsilon)     AQKFNGLTVLPPLLTDEMIAQYTSALLAGTITSGWTFGAGAALQIPFAMQMAYRFNGIGV    911
B.1.525(Eta)         AQKFNGLTVLPPLLTDEMIAQYTSALLAGTITSGWTLGAGAALQIPFAMQMAYRFNGIGV    908
B.1.526(Iota)        AQKFNGLTVLPPLLTDEMIAQYTSALLAGTITSGWTFGAGAALQIPFAMQMAYRFNGIGV    911
B.1.526.1            AQKFNGLNVLPPLLTDEMIAQYTSALLAGTITSGWTFGAGAALQIPFAMQMAYRFNGIGV    910
B.1.617.1(Kappa)     AQKFNGLTVLPPLLTDEMIAQYTSALLAGTITSGWTFGAGAALQIPFAMQMAYRFNGIGV    911
B.1.617.2(Delta)     AQKFNGLTVLPPLLTDEMIAQYTSALLAGTITSGWTFGAGAALQIPFAMQMAYRFNGIGV    909
P.1(Gamma)           AQKFNGLTVLPPLLTDEMIAQYTSALLAGTITSGWTFGAGAALQIPFAMQMAYRFNGIGV    911
P.2(Zeta)            AQKFNGLTVLPPLLTDEMIAQYTSALLAGTITSGWTFGAGAALQIPFAMQMAYRFNGIGV    911
B.1.1.529            AQKFKGLTVLPPLLTDEMIAQYTSALLAGTITSGWTFGAGAALQIPFAMQMAYRFNGIGV    908
                     ** . * .*****:  :* * **::*:..    *** * .:   **** .: **:**:*:

                                                         950 954             969
MERS                 TQQVLSENQKLIANKFNQALGAMQTGFTTTNEAFQKVQDAVNNNAQALSKLASELSNTFG    1045
SARS                 TQNVLYENQKQIANQFNKAISQIQESLTTTSTALGKLQDVVNQNAQALNTLVKQLSSNFG    953
WA1(COVID-19)        TQNVLYENQKLIANQFNSAIGKIQDSLSSTASALGKLQDVVNQNAQALNTLVKQLSSNFG    971
B.1.17(Alpha)        TQNVLYENQKLIANQFNSAIGKIQDSLSSTASALGKLQDVVNQNAQALNTLVKQLSSNFG    968
B.1.351(Beta)        TQNVLYENQKLIANQFNSAIGKIQDSLSSTASALGKLQDVVNQNAQALNTLVKQLSSNFG    968
B.1.427(Epsilon)     TQNVLYENQKLIANQFNSAIGKIQDSLSSTASALGKLQDVVNQNAQALNTLVKQLSSNFG    971
B.1.429(Epsilon)     TQNVLYENQKLIANQFNSAIGKIQDSLSSTASALGKLQDVVNQNAQALNTLVKQLSSNFG    971
B.1.525(Eta)         TQNVLYENQKLIANQFNSAIGKIQDSLSSTASALGKLQDVVNQNAQALNTLVKQLSSNFG    968
B.1.526(Iota)        TQNVLYENQKLIANQFNSAIGKIQDSLSSTASALGKLQDVVNQNAQALNTLVKQLSSNFG    971
B.1.526.1            TQNVLYENQKLIANQFNSAIGKIQDSLSSTASALGKLQHVVNQNAQALNTLVKQLSSNFG    970
B.1.617.1(Kappa)     TQNVLYENQKLIANQFNSAIGKIQDSLSSTASALGKLQDVVNQNAQALNTLVKQLSSNFG    971
B.1.617.2(Delta)     TQNVLYENQKLIANQFNSAIGKIQDSLSSTASALGKLQNVVNQNAQALNTLVKQLSSNFG    969
P.1(Gamma)           TQNVLYENQKLIANQFNSAIGKIQDSLSSTASALGKLQDVVNQNAQALNTLVKQLSSNFG    971
P.2(Zeta)            TQNVLYENQKLIANQFNSAIGKIQDSLSSTASALGKLQDVVNQNAQALNTLVKQLSSNFG    971
B.1.1.529            TQNVLYENQKLIANQFNSAIGKIQDSLSSTASALGKLQDVVNHNAQALNTLVKQLSSKFG    968
                     **:** **** ***:**.*:. :* .:::*  *: *:*..**:*****..*..:**..**

                          981 982                                       1027
MERS                 AISASIGDIIQRLDVLEQDAQIDRLINGRLTTLNAFVAQQLVRSESAALSAQLAKDKVNE    1105
SARS                 AISSVLNDILSRLDKVEAEVQIDRLITGRLQSLQTYVTQQLIRAAEIRASANLAATKMSE    1013
WA1(COVID-19)        AISSVLNDILSRLDKVEAEVQIDRLITGRLQSLQTYVTQQLIRAAEIRASANLAATKMSE    1031
B.1.17(Alpha)        AISSVLNDILARLDKVEAEVQIDRLITGRLQSLQTYVTQQLIRAAEIRASANLAATKMSE    1028
B.1.351(Beta)        AISSVLNDILSRLDKVEAEVQIDRLITGRLQSLQTYVTQQLIRAAEIRASANLAATKMSE    1028
B.1.427(Epsilon)     AISSVLNDILSRLDKVEAEVQIDRLITGRLQSLQTYVTQQLIRAAEIRASANLAATKMSE    1031
B.1.429(Epsilon)     AISSVLNDILSRLDKVEAEVQIDRLITGRLQSLQTYVTQQLIRAAEIRASANLAATKMSE    1031
B.1.525(Eta)         AISSVLNDILSRLDKVEAEVQIDRLITGRLQSLQTYVTQQLIRAAEIRASANLAATKMSE    1028
B.1.526(Iota)        AISSVLNDILSRLDKVEAEVQIDRLITGRLQSLQTYVTQQLIRAAEIRASANLAATKMSE    1031
B.1.526.1            AISSVLNDILSRLDKVEAEVQIDRLITGRLQSLQTYVTQQLIRAAEIRASANLAATKMSE    1030
B.1.617.1(Kappa)     AISSVLNDILSRLDKVEAEVQIDRLITGRLQSLQTYVTQQLIRAAEIRASANLAATKMSE    1031
B.1.617.2(Delta)     AISSVLNDILSRLDKVEAEVQIDRLITGRLQSLQTYVTQQLIRAAEIRASANLAATKMSE    1029
P.1(Gamma)           AISSVLNDILSRLDKVEAEVQIDRLITGRLQSLQTYVTQQLIRAAEIRASANLAAIKMSE    1031
P.2(Zeta)            AISSVLNDILSRLDKVEAEVQIDRLITGRLQSLQTYVTQQLIRAAEIRASANLAATKMSE    1031
B.1.1.529            AISSVLNDIFSRLDKVEAEVQIDRLITGRLQSLQTYVTQQLIRAAEIRASANLAATKMSE    1028
                     ***: :.**: *** :* :.******.*** :*:::*:***:*: .   **:**  *:.*

                                                         1071
MERS                 CVKAQSKRSGFCGQGTHIVSFVVNAPNGLYFMHVGYYPSNHIEVVSAYGLCDAANPTNCI    1165
SARS                 CVLGQSKRVDFCGKGYHLMSFPQAAPHGVVFLHVTYVPSQERNFTTAPAICHEGKA---Y    1070
WA1(COVID-19)        CVLGQSKRVDFCGKGYHLMSFPQSAPHGVVFLHVTYVPAQEKNFTTAPAICHDGKA---H    1088
B.1.17(Alpha)        CVLGQSKRVDFCGKGYHLMSFPQSAPHGVVFLHVTYVPAQEKNFTTAPAICHDGKA---H    1085
B.1.351(Beta)        CVLGQSKRVDFCGKGYHLMSFPQSAPHGVVFLHVTYVPAQEKNFTTAPAICHDGKA---H    1085
B.1.427(Epsilon)     CVLGQSKRVDFCGKGYHLMSFPQSAPHGVVFLHVTYVPAQEKNFTTAPAICHDGKA---H    1088
B.1.429(Epsilon)     CVLGQSKRVDFCGKGYHLMSFPQSAPHGVVFLHVTYVPAQEKNFTTAPAICHDGKA---H    1088
B.1.525(Eta)         CVLGQSKRVDFCGKGYHLMSFPQSAPHGVVFLHVTYVPAQEKNFTTAPAICHDGKA---H    1085
B.1.526(Iota)        CVLGQSKRVDFCGKGYHLMSFPQSAPHGVVFLHVTYVPAQEKNFTTAPAICHDGKA---H    1088
B.1.526.1            CVLGQSKRVDFCGKGYHLMSFPQSAPHGVVFLHVTYVPAQEKNFTTAPAICHDGKA---H    1087
B.1.617.1(Kappa)     CVLGQSKRVDFCGKGYHLMSFPQSAPHGVVFLHVTYVPAHEKNFTTAPAICHDGKA---H    1088
B.1.617.2(Delta)     CVLGQSKRVDFCGKGYHLMSFPQSAPHGVVFLHVTYVPAQEKNFTTAPAICHDGKA---H    1086
P.1(Gamma)           CVLGQSKRVDFCGKGYHLMSFPQSAPHGVVFLHVTYVPAQEKNFTTAPAICHDGKA---H    1088
P.2(Zeta)            CVLGQSKRVDFCGKGYHLMSFPQSAPHGVVFLHVTYVPAQEKNFTTAPAICHDGKA---H    1088
B.1.1.529            CVLGQSKRVDFCGKGYHLMSFPQSAPHGVVFLHVTYVPAQEKNFTTAPAICHDGKA---H    1085
                     ** .**** .***:* *::**   **:*: *:** * *::. :..:* .:*. .:
```

```
                                      1101                         1118
MERS                APVNGYFIKTNNTRIVDEWSYTGSSFYAPEPITSLNTKYVAPQVTYQN-ISTNLPPPLLG    1224
SARS                FPREGVFVFN-----GTSWFITQRNFFSPQIITTDNTFVSGNCDVVIGIINNTVYDPLQ-    1124
WA1(COVID-19)       FPREGVFVSN-----GTHWFVTQRNFYEPQIITTDNTFVSGNCDVVIGIVNNTVYDPLQ-    1142
B.1.17(Alpha)       FPREGVFVSN-----GTHWFVTQRNFYEPQIITTHNTFVSGNCDVVIGIVNNTVYDPLQ-    1139
B.1.351(Beta)       FPREGVFVSN-----GTHWFVTQRNFYEPQIITTDNTFVSGNCDVVIGIVNNTVYDPLQ-    1139
B.1.427(Epsilon)    FPREGVFVSN-----GTHWFVTQRNFYEPQIITTDNTFVSGNCDVVIGIVNNTVYDPLQ-    1142
B.1.429(Epsilon)    FPREGVFVSN-----GTHWFVTQRNFYEPQIITTDNTFVSGNCDVVIGIVNNTVYDPLQ-    1142
B.1.525(Eta)        FPREGVFVSN-----GTHWFVTQRNFYEPQIITTDNTFVSGNCDVVIGIVNNTVYDPLQ-    1139
B.1.526(Iota)       FPREGVFVSN-----GTHWFVTQRNFYEPQIITTDNTFVSGNCDVVIGIVNNTVYDPLQ-    1142
B.1.526.1           FPREGVFVSN-----GTHWFVTQRNFYEPQIITTDNTFVSGNCDVVIGIVNNTVYDPLQ-    1141
B.1.617.1(Kappa)    FPREGVFVSN-----GTDWFVTQRNFYEPQIITTDNTFVSGNCDVVIGIVNNTVYDPLQ-    1142
B.1.617.2(Delta)    FPREGVFVSN-----GTHWFVTQRNFYEPQIITTDNTFVSGNCDVVIGIVNNTVYDPLQ-    1140
P.1(Gamma)          FPREGVFVSN-----GTHWFVTQRNFYEPQIITTDNTFVSGNCDVVIGIVNNTVYDPLQ-    1142
P.2(Zeta)           FPREGVFVSN-----GTHWFVTQRNFYEPQIITTDNTFVSGNCDVVIGIVNNTVYDPLQ-    1142
B.1.1.529           FPREGVFVSN-----GTHWFVTQRNFYEPQIITTDNTFVSGNCDVVIGIVNNTVYDPLQ-    1139
                     * :* *: .         *  *  .*: *: **: **    .    .  . :...:  **

                                                   1176
MERS                NSTGIDFQDELDEFFKNVSTSIPNFGSLTQINTTLLDLTYEMLSLQQVVKALNESYIDLK    1284
SARS                -PELDSFKEELDKYFKNHTSPDVDLGDISGINASVVNIQKEIDRLNEVAKNLNESLIDLQ    1183
WA1(COVID-19)       -PELDSFKEELDKYFKNHTSPDVDLGDISGINASVVNIQKEIDRLNEVAKNLNESLIDLQ    1201
B.1.17(Alpha)       -PELDSFKEELDKYFKNHTSPDVDLGDISGINASVVNIQKEIDRLNEVAKNLNESLIDLQ    1198
B.1.351(Beta)       -PELDSFKEELDKYFKNHTSPDVDLGDISGINASVVNIQKEIDRLNEVAKNLNESLIDLQ    1198
B.1.427(Epsilon)    -PELDSFKEELDKYFKNHTSPDVDLGDISGINASVVNIQKEIDRLNEVAKNLNESLIDLQ    1201
B.1.429(Epsilon)    -PELDSFKEELDKYFKNHTSPDVDLGDISGINASVVNIQKEIDRLNEVAKNLNESLIDLQ    1201
B.1.525(Eta)        -PELDSFKEELDKYFKNHTSPDVDLGDISGINASVVNIQKEIDRLNEVAKNLNESLIDLQ    1198
B.1.526(Iota)       -PELDSFKEELDKYFKNHTSPDVDLGDISGINASVVNIQKEIDRLNEVAKNLNESLIDLQ    1201
B.1.526.1           -PELDSFKEELDKYFKNHTSPDVDLGDISGINASVVNIQKEIDRLNEVAKNLNESLIDLQ    1200
B.1.617.1(Kappa)    -PELDSFKEELDKYFKNHTSPDVDLGDISGINASVVNIQKEIDRLNEVAKNLNESLIDLQ    1201
B.1.617.2(Delta)    -PELDSFKEELDKYFKNHTSPDVDLGDISGINASVVNIQKEIDRLNEVAKNLNESLIDLQ    1199
P.1(Gamma)          -PELDSFKEELDKYFKNHTSPDVDLGDISGINASFVNIQKEIDRLNEVAKNLNESLIDLQ    1201
P.2(Zeta)           -PELDSFKEELDKYFKNHTSPDVDLGDISGINASFVNIQKEIDRLNEVAKNLNESLIDLQ    1201
B.1.1.529           -PELDSFKEELDKYFKNHTSPDVDLGDISGINASVVNIQKEIDRLNEVAKNLNESLIDLQ    1198
                         .*::***::*** ::   ::*.:: **::.:::  *:   *::*.* **** ***:


MERS                ELGNYTYYNKWPWYIWLGFIAGLVALALCVFFILCCTGCGTNCMGKLKCNRCCDRYEEYD    1344
SARS                ELGKYEQYIKWPWYVWLGFIAGLIAIVMVTILLCCMTSCCSCLKGACSCGSCCKF-DEDD    1242
WA1(COVID-19)       ELGKYEQYIKWPWYIWLGFIAGLIAIVMVTIMLCCMTSCCSCLKGCCSCGSCCKF-DEDD    1260
B.1.17(Alpha)       ELGKYEQYIKWPWYIWLGFIAGLIAIVMVTIMLCCMTSCCSCLKGCCSCGSCCKF-DEDD    1257
B.1.351(Beta)       ELGKYEQYIKWPWYIWLGFIAGLIAIVMVTIMLCCMTSCCSCLKGCCSCGSCCKF-DEDD    1257
B.1.427(Epsilon)    ELGKYEQYIKWPWYIWLGFIAGLIAIVMVTIMLCCMTSCCSCLKGCCSCGSCCKF-DEDD    1260
B.1.429(Epsilon)    ELGKYEQYIKWPWYIWLGFIAGLIAIVMVTIMLCCMTSCCSCLKGCCSCGSCCKF-DEDD    1260
B.1.525(Eta)        ELGKYEQYIKWPWYIWLGFIAGLIAIVMVTIMLCCMTSCCSCLKGCCSCGSCCKF-DEDD    1257
B.1.526(Iota)       ELGKYEQYIKWPWYIWLGFIAGLIAIVMVTIMLCCMTSCCSCLKGCCSCGSCCKF-DEDD    1260
B.1.526.1           ELGKYEQYIKWPWYIWLGFIAGLIAIVMVTIMLCCMTSCCSCLKGCCSCGSCCKF-DEDD    1259
B.1.617.1(Kappa)    ELGKYEQYIKWPWYIWLGFIAGLIAIVMVTIMLCCMTSCCSCLKGCCSCGSCCKF-DEDD    1260
B.1.617.2(Delta)    ELGKYEQYIKWPWYIWLGFIAGLIAIVMVTIMLCCMTSCCSCLKGCCSCGSCCKF-DEDD    1258
P.1(Gamma)          ELGKYEQYIKWPWYIWLGFIAGLIAIVMVTIMLCCMTSCCSCLKGCCSCGSCCKF-DEDD    1260
P.2(Zeta)           ELGKYEQYIKWPWYIWLGFIAGLIAIVMVTIMLCCMTSCCSCLKGCCSCGSCCKF-DEDD    1260
B.1.1.529           ELGKYEQYIKWPWYIWLGFIAGLIAIVMVTIMLCCMTSCCSCLKGCCSCGSCCKF-DEDD    1257
                    ***:*   * *****:********:*:.: .::: * *.* :    *   .*. **.  :* *

MERS                LEPHKVHVH----     1353
SARS                SEPVLKGVKLHYT     1255
WA1(COVID-19)       SEPVLKGVKLHYT     1273
B.1.17(Alpha)       SEPVLKGVKLHYT     1270
B.1.351(Beta)       SEPVLKGVKLHYT     1270
B.1.427(Epsilon)    SEPVLKGVKLHYT     1273
B.1.429(Epsilon)    SEPVLKGVKLHYT     1273
B.1.525(Eta)        SEPVLKGVKLHYT     1270
B.1.526(Iota)       SEPVLKGVKLHYT     1273
B.1.526.1           SEPVLKGVKLHYT     1272
B.1.617.1(Kappa)    SEPVLKGVKLHYT     1273
B.1.617.2(Delta)    SEPVLKGVKLHYT     1271
P.1(Gamma)          SEPVLKGVKLHYT     1273
P.2(Zeta)           SEPVLKGVKLHYT     1273
B.1.1.529           SEPVLKGVKLHYT     1270
                     **    *:
```

**Supplementary Figure S2**
